# Atherosclerosis destabilizes regulatory T cells (Tregs) resulting in multiple families of exTregs

**DOI:** 10.64898/2026.07.25.740705

**Authors:** Qingkang Lyu, Yan Wang, Anusha Bellapu, Polina Bombina, Sunil Kumar, Lauren Fogel, Smriti Parashar, Kunzhe Dong, Jiang Zhou, Devadatta Ashok Gosavi, Mikhail Fomin, Klaus Ley

**Author notes:** Corresponding author: Klaus Ley, Immunology Center of Georgia, Augusta University, Augusta, GA 30912, USA; Email.

## Abstract

How regulatory T cells (Tregs) lose lineage identity during chronic inflammation remains poorly understood. Here, using inducible Foxp3 lineage tracing together with single-cell transcriptomic, proteomic and T cell receptor (TCR) profiling in atherosclerosis-prone mice, we identify Treg destabilization as a staged and branching differentiation process rather than an abrupt loss of lineage identity. Conventional Tregs (cTregs) first transition through an effector Treg (eTreg) intermediate characterized by attenuation of the CD25–STAT5 axis while retaining core Treg features, before diversifying into eight transcriptionally distinct exTreg states, including Tfh-like, cytotoxic, Th1-like inflammatory, Th1-like cytotoxic and proliferative populations. Trajectory inference, TCR clonotype analysis and experimental Treg-to-exTreg conversion independently converged on this developmental framework, revealing that clonally related exTregs acquire distinct effector programs. Mechanistically, we identify Treg-intrinsic IL-6R signaling as an important driver of this process. IL-6 accelerated exTreg generation in vitro, whereas Treg-specific deletion of Il6ra reduced inflammatory exTreg differentiation and attenuated atherosclerosis in vivo. Together, these findings establish a framework for Treg destabilization during atherosclerosis and provide a conceptual basis for preserving Treg lineage stability in chronic inflammatory disease.

## Introduction

Atherosclerosis is a chronic inflammatory disease of the arterial wall characterized by sustained immune activation and disruption of immune homeostasis ^1, 2^. In addition to metabolic and environmental risk factors, accumulating evidence indicates that immune-mediated mechanisms, including features of autoimmunity and loss of immune tolerance, critically shape disease initiation and progression ^3, 4^. One known self-antigen in atherosclerosis is apolipoprotein B (ApoB), the core lipoprotein of low-density lipoprotein (LDL) and chylomicrons. ApoB-specific CD4⁺ T cells are detectable in both mice and humans and expand with disease progression ^5^. While ApoB-reactive T cells exhibit regulatory features under homeostatic conditions ^6^, advancing atherosclerosis is associated with their progressive acquisition of effector-like programs, accompanied by attenuation of canonical Treg gene expression ^5, 7, 8^.

Regulatory T (Treg) cells, a specialized CD4⁺ T cell lineage defined by expression of the transcription factor Foxp3, are essential for maintaining self-tolerance and restraining pathologic autoimmune responses. Consistent with this role, experimental studies have shown that impairment of Treg function exacerbates atherosclerosis, whereas augmentation of Treg activity limits disease progression, supporting a central role for Treg cells in controlling chronic vascular inflammation ^9, 10, 11, 12^. Notably, Treg cells within atherosclerotic tissues exhibit dynamic regulation ^13^ and mediate protection in part through immunosuppressive cytokines ^14, 15, 16, 17^, underscoring their functional relevance within the inflamed vascular microenvironment.

Although Treg cells exert protective effects in atherosclerosis, accumulating evidence suggests that Treg cells are not a static population but display phenotypic and functional heterogeneity in inflammatory environments ^17^.

Under physiological conditions, Treg cells maintain high expression of Foxp3 and the interleukin-2 receptor α-chain (CD25), enforcing a transcriptional and epigenetic program that supports suppressive function and immune homeostasis. To fulfill their regulatory roles across diverse tissue niches and immune contexts, Treg cells display a degree of functional plasticity that enables adaptation to local cues ^18, 19, 20^. This plasticity includes the acquisition of features associated with conventional T helper cell lineages, such as expression of T-bet for T-helper (Th1), RORγt for Th17, or GATA3 for Th2, while retaining Foxp3 expression and suppressive capacity. For example, during type 1 inflammatory responses, regulatory T cells can upregulate the Th1-associated transcription factor T-bet and the chemokine receptor CXCR3, enabling their migration to and selective suppression of Th1-polarized inflammatory sites ^21, 22, 23^. Genetic fate-mapping studies have demonstrated that T-bet-expressing Treg cells represent a stable and functionally specialized subset that retains Foxp3 expression and suppressive capacity in healthy mice, rather than undergoing conversion into effector Th1 cells ^24^. Such “Th-like” Treg states are able to prevent or curb autoimmunity by enhancing context-specific immune regulation and restraining polarized effector responses.

The same adaptive flexibility that supports context-specific Treg function may compromise lineage stability under conditions of chronic inflammation, metabolic stress, or cytokine imbalance. In these settings, Treg cells can downregulate Foxp3, lose suppressive activity, and acquire pro-inflammatory or cytotoxic effector programs, giving rise to cells commonly referred to as exTregs^25, 26^. These unstable Treg-derived cells frequently exhibit reduced or absent CD25 expression, acquisition of lineage-defining transcription factors (such as T-bet and RORγt), production of inflammatory cytokines such as IFN-γ or IL-17 ^27, 28^, and epigenetic remodeling at conserved regulatory elements of the Foxp3 locus ^29^. exTreg populations have been reported across multiple autoimmune and inflammatory diseases such as rheumatoid arthritis, multiple sclerosis, atherosclerosis, allograft rejection, and viral infection, implicating Treg instability as a potential driver of immune dysregulation ^7, 8, 17, 28, 30, 31^.

Evidence for Treg destabilization is particularly relevant to atherosclerosis. Lineage-tracing studies have shown that TFH-like exTregs can arise from bona fide Foxp3+ Tregs and lose their atheroprotective capacity as inflammatory burden increases ^32^. In human coronary artery disease, ApoB-reactive Tregs occupy an effector Treg state characterized by tissue-homing receptors and plaque tropism. These cells are antigen-driven and clonally expanded and, with increasing inflammatory burden, downregulate Treg lineage genes while acquiring pro-inflammatory and cytotoxic programs consistent with exTreg-like conversion ^33^. Recent studies further illustrate the diversity of exTreg phenotypes: cytotoxic CD4+ exTregs have been identified in human blood, whereas Th1-like, Th17-like and TFH-like exTreg populations have been described in inflammatory and autoimmune settings ^7, 8, 28, 32^

Despite growing evidence for Treg instability in atherosclerosis, how stable Tregs progress toward exTreg states remains poorly understood. In particular, it is unclear whether loss of Treg identity occurs as an abrupt lineage switch or through defined intermediate states. It is also unknown whether distinct exTreg phenotypes arise independently or through coordinated differentiation trajectories. The inflammatory signals that promote this process also remain incompletely defined. IL-6 is a candidate regulator of Treg instability because IL-6–IL-6R signaling can antagonize Foxp3 maintenance through STAT3-dependent mechanisms and has been linked to Treg-to-TFH conversion during atherosclerosis ^27, 32, 34^. However, whether Treg-intrinsic IL-6R signaling promotes broader Treg-to-exTreg conversion and diversification during atherosclerosis remains unknown.

Here, we used Foxp3 lineage-tracing mice on the atherosclerosis-prone *Apoe*^-/-^background together with single-cell RNA sequencing, CITE-seq and TCR sequencing to resolve Treg-lineage states during atherosclerosis. We identify an effector Treg state that bridges conventional Tregs and exTregs and show that exTregs subsequently diversify along distinct, clonally connected differentiation trajectories. We further identify Treg-intrinsic IL-6R signaling as a promoter of Treg-to-exTreg conversion and show that its disruption reduces inflammatory exTreg states and atherosclerosis. Together, our findings define Treg destabilization as a staged and organized differentiation process rather than an abrupt loss of Treg identity.

## Results

### Progressive loss of IL-2 responsiveness accompanies Treg destabilization during atherosclerosis

To investigate how Tregs progressively lose lineage stability during atherosclerosis, we established an inducible FoxP3 lineage-tracing *Apoe^-/-^* mouse model in which mice received two rounds of tamoxifen administration before and during 12 weeks of Western diet feeding to permanently label cells with a history of Foxp3 expression (Fig. 1a). As expected, Western diet feeding induced robust atherosclerosis, evidenced by significantly increased plaque area and plaque burden throughout the aorta compared with chow-fed controls (Extended Data Fig. 1a–c). Concomitantly, Western diet-fed mice exhibited expansion of splenic and lymph node CD4+ T cells together with increased frequencies of both Tregs (Foxp3-GFP+tdTomato+) and exTregs (Foxp3-GFP-tdTomato+), indicating that chronic hyperlipidemia promotes Treg activation and destabilization (Extended Data Fig. 1d, e). To investigate the molecular basis of Treg destabilization, splenic CD4+ T cells from Foxp3 lineage-tracing mice were analyzed by flow cytometry and subjected to CITE-seq labeling followed by single-cell RNA and TCR sequencing (Fig. 1b).

**Fig. 1.**
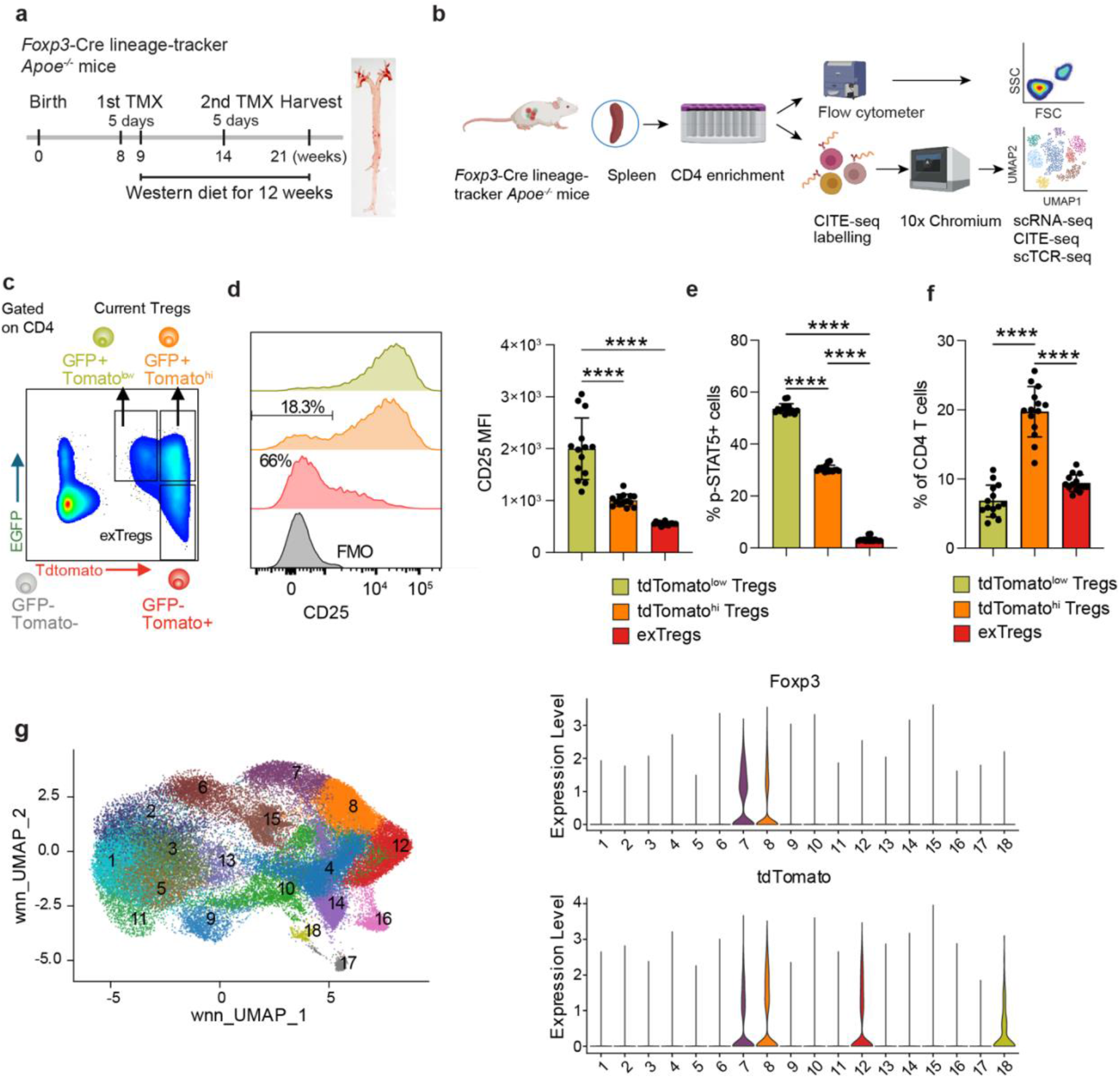
Progressive loss of IL-2 responsiveness accompanies Treg destabilization during atherosclerosis. **a,** Experimental design of the Foxp3 lineage-tracing Apoe^-/-^ mouse model. Mice received two rounds of tamoxifen administration before and during Western diet feeding to permanently label cells with a history of Foxp3 expression and were analyzed after 12 weeks of Western diet feeding. **b,** Experimental workflow for flow cytometry and integrated single-cell analyses. Splenic CD4+ T cells isolated from Foxp3 lineage-tracing Apoe^-/-^ mice were analyzed by flow cytometry or subjected to CITE-seq labeling followed by 10x Genomics single-cell RNA sequencing and paired TCR sequencing. **c,** Representative flow cytometric gating strategy identifying tdTomato^low^ Tregs, tdTomato^hi^ Tregs and GFP-tdTomato+ exTregs among lineage-traced CD4+ T cells. GFP identifies current Foxp3-expressing Tregs, whereas GFP-tdTomato+ cells represent exTregs. **d,** Representative histograms (left) and quantification of CD25 mean fluorescence intensity (MFI, right) in tdTomato^low^ Tregs, tdTomato^hi^ Tregs and exTregs. **e,** Enriched CD4+ T cells were stimulated ex vivo with recombinant IL-2 (500 U/ml, 20 min), followed by intracellular staining for phosphorylated STAT5 (pSTAT5) and flow cytometric analysis. Quantification of pSTAT5+ cells in tdTomato^low^ Tregs, tdTomato^hi^ Tregs and exTregs. **f,** Frequencies of tdTomato^low^ Tregs, tdTomato^hi^ Tregs and exTregs among splenic CD4+ T cells. **g,** Weighted nearest-neighbor (WNN) analysis of the integrated single-cell dataset showing 18 CD4+ T-cell clusters (left). Violin plots showing Foxp3 (top) and tdTomato (bottom) expression across individual clusters (right). Data are presented as mean ± SEM. Each dot represents one biological replicate (mouse). Statistical significance was determined using one-way ANOVA followed by Tukey’s multiple-comparison test unless otherwise indicated. \**P* < 0.05, \*\**P* < 0.01, \*\*\**P* < 0.001, **\*\*\*\****P* < 0.0001.

Flow cytometric analysis consistently resolved three FoxP3-lineage Treg states: GFP+tdTomato^low^ Tregs, GFP+tdTomato^hi^ Tregs, and GFP-tdTomato+ exTregs (Fig. 1c). Because IL-2 signaling is essential for maintaining Treg identity, we next examined CD25 expression across these three populations. CD25 expression was uniformly high in tdTomato^low^ Tregs, began to decline in tdTomato^hi^ Tregs, and was markedly reduced in exTregs (Fig. 1d). Correspondingly, phosphorylation of STAT5 was progressively reduced from tdTomato^low^ Tregs to tdTomato^hi^ Tregs to exTregs (Fig. 1e), indicating gradual attenuation of IL-2 responsiveness. Notably, tdTomato^hi^ Tregs constituted the predominant Treg population in the spleen (Fig. 1f). These observations identified tdTomato^hi^ Tregs as the predominant Treg population with attenuated IL-2 responsiveness and prompted us to define their molecular identity and relationship to exTregs.

Integrated single-cell multiomic profiling (Fig. 1b) by weighted nearest-neighbor^35^ analysis identified 18 transcriptionally distinct CD4+ T-cell clusters (Fig. 1g). Foxp3 expression was largely restricted to clusters 7 and 8, whereas tdTomato expression was enriched in clusters 7, 8, 12 and 18, revealing substantial heterogeneity among Foxp3 lineage-traced CD4+ T cells.

### Treg destabilization proceeds through a staged cTreg-to-eTreg-to-exTreg transition

Next, we investigated the progression of Treg destabilization using integrated single-cell multiomic profiling. As observed by flow cytometry, tdTomato expression displayed a clear bimodal distribution in the single-cell dataset. We applied ThresholdR^36^ to objectively determine the optimal cutoff separating tdTomato^low^ and tdTomato^hi^ populations (Extended Data 2a). This identified resting cTregs (yellow in Fig. 2a), effector eTregs (orange in Fig. 2a) and exTregs (red in Fig. 2a). Density-based mapping of GFP-tdTomato+ cells revealed that exTregs were highly enriched within a discrete region of the integrated UMAP and sparsely distributed elsewhere (Extended Data 2b). The exTreg region was CD44^hi^, whereas Tregs were CD25^hi^ and CD62L^hi^ (Extended Data 2c).

**Fig. 2.**
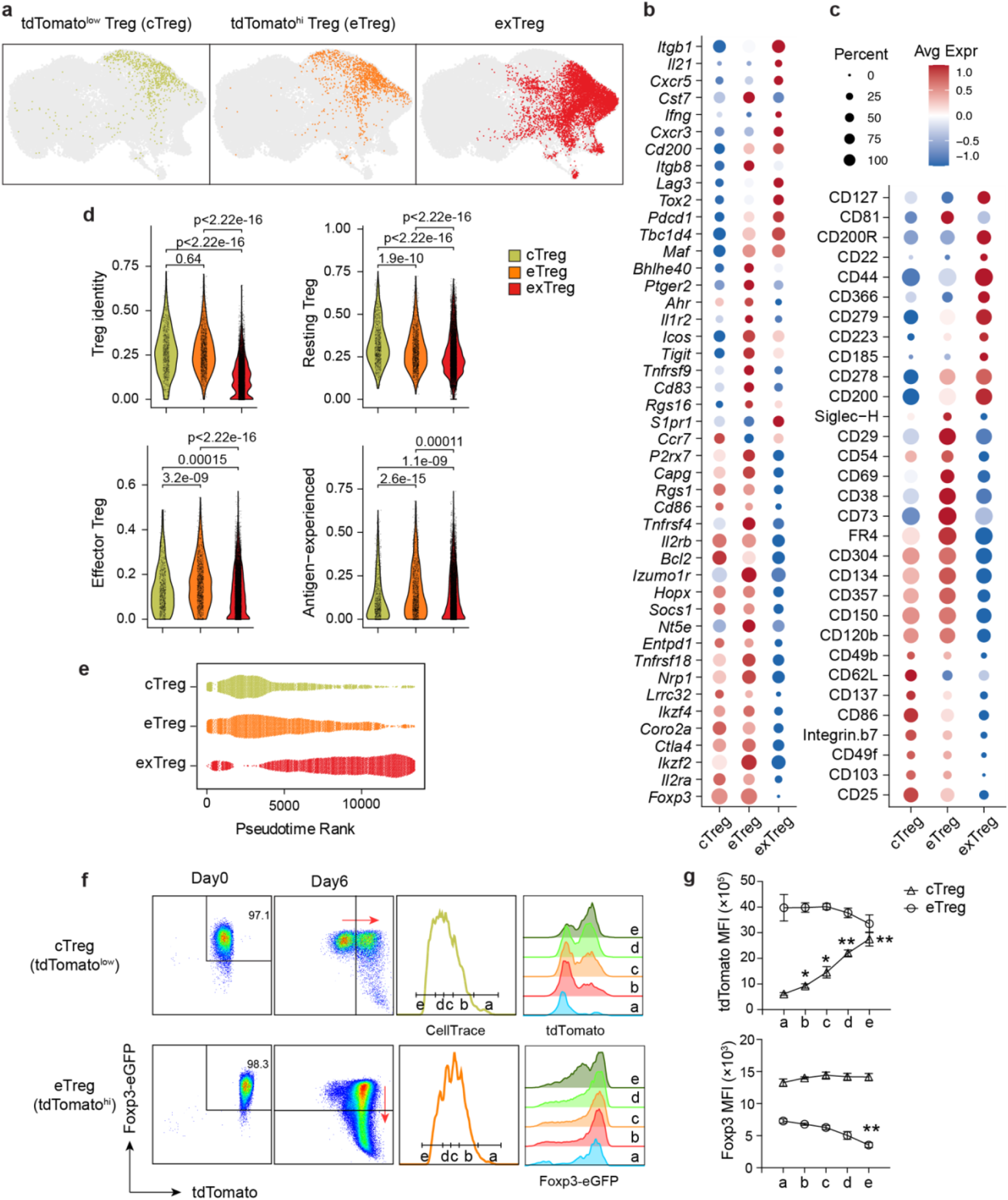
Stepwise conversion of Tregs into exTregs through an effector Treg intermediate. **a,** UMAP visualization of integrated scRNA-seq/CITE-seq data showing the distribution of Foxp3+tdTomato^low^ Tregs, Foxp3+tdTomato^hi^ Tregs, and GFP-tdTomato+ exTregs identified from Foxp3 lineage-tracing Apoe^−/−^ mice. **b,** Dot plot showing the top 15 differentially expressed genes (DEGs) identified by scRNA-seq for each Treg population. **c,** Dot plot showing the top 10 differentially abundant surface proteins identified by CITE-seq for each Treg population. **d,** AUCell module score analysis using previously published transcriptional signatures representing Treg identity, resting Tregs, effector Tregs and antigen-experienced T cells. AUCell scores were calculated from single-cell RNA-sequencing data using previously defined gene modules. Each dot represents one cell. Violin widths indicate the distribution of AUCell scores within each population. Statistical significance was determined using a two-sided Wilcoxon rank-sum test, and adjusted P values are shown. **e,** Monocle3 pseudotime ranking showing the relative positions of cTregs, eTregs and exTregs along the inferred differentiation trajectory. **f,** In vitro differentiation assay. FACS-sorted tdTomato^low^ Tregs (cTregs) and tdTomato^hi^ Tregs (eTregs) were labeled with CellTrace Violet (CTV) and cultured for 6 days in the presence of anti-CD3/CD28 antibody-coated beads and recombinant IL-2 (200 U/ml). Representative flow cytometry plots show the transition of cTregs into tdTomato^hi^ Tregs while maintaining Foxp3 expression, whereas eTregs progressively lost Foxp3 and converted into GFP-tdTomato+ exTregs. Cells were further stratified into five successive generations according to CTV dilution (a–e) to examine tdTomato and Foxp3 expression during cell division. **g**, Representative histograms and quantification from three independent experiments show progressive upregulation of tdTomato expression across successive generations in cTregs, whereas Foxp3 expression progressively declined in eTregs. Unless otherwise indicated, all analyses were performed using splenic CD4+ T cells isolated from Foxp3 lineage-tracing *Apoe*^-/-^mice. Data are presented as mean ± SEM. Each dot represents one biological replicate. Statistical significance was determined by one-way ANOVA with Tukey’s multiple-comparison test, unless otherwise indicated. \**P* < 0.05, \*\**P* < 0.01, \*\*\**P* < 0.001, **\*\*\*\****P* < 0.0001.

Transcriptomic and proteomic profiling demonstrated that these three populations represented distinct molecular states (Fig. 2b, c). tdTomato^low^ Tregs expressed a canonical Treg program characterized by high expression of *Foxp3*, *Il2ra*, *Ikzf2*, *Ikzf4*, *Nrp1* and other lineage-associated genes together with high surface expression of CD25, CD103, CD137, CD304 and CD357. Notably, tdTomato^hi^ Tregs retained core Treg lineage markers but displayed reduced Il2ra (encoding CD25) expression together with increased expression of activation-associated genes including *Icos*, *Tigit*, *Tnfrsf4* and *Tnfrsf18*, accompanied by increased expression of effector-associated surface molecules such as CD38, CD69, CD73, PD-1 (CD279) and CD29 (encoded by Itgb1). In contrast, exTregs acquired inflammatory and effector-associated features, including increased expression of *Itgb1, Il21*, *Cxcr5*, *Cxcr3*, *Ifng*, *Lag3*, *Tox2*, *Pdcd1*, *Maf* and multiple activation-associated surface proteins. Together, these molecular features indicate that eTregs retain core Treg identity while acquiring effector-associated characteristics.

Gene signature^37, 38, 39, 40^ analysis further supported the identities of these populations (Fig. 2d). Treg identity and resting Treg signatures progressively declined from tdTomato^low^ Tregs to tdTomato^hi^ Tregs and reached their lowest levels in exTregs. The effector Treg signature was strongest in eTregs. exTregs were enriched in an antigen-experienced signature (Fig. 2d). Together, these analyses identify tdTomato^low^ and tdTomato^hi^ Tregs as conventional Tregs (cTregs) and effector Tregs (eTregs), respectively, positioning eTregs between cTregs and exTregs.

Monocle3 trajectory inference positioned cTregs at the beginning of the differentiation path, eTregs at an intermediate position and exTregs at the terminal state (Extended Data Fig. 3a). Pseudotime ordering further supported progressive differentiation from cTregs through eTregs to exTregs (Fig. 2e). To experimentally validate this differentiation pathway, we sorted cTregs and eTregs and cultured them for six days with anti-CD3/CD28 beads in the presence of IL-2. Most cTregs acquired increased tdTomato expression while maintaining Foxp3, consistent with differentiation into eTregs, whereas a smaller fraction lost Foxp3 and became exTregs (Fig. 2f). In contrast, eTregs efficiently generated exTregs but did not revert to cTregs. CellTrace Violet (CTV) dilution demonstrated that tdTomato expression progressively increased with successive cell divisions in cTregs, whereas Foxp3 expression progressively declined during proliferation of eTregs (Fig. 2f). A gradual rise of tdTomato expression mirrored progressive loss of FoxP3 (Fig. 2g). Together, these computational and functional analyses support a staged model of Treg destabilization in which conventional Tregs first acquire an effector Treg state before progressively differentiating into exTregs.

### Treg destabilization generates diverse effector exTreg states

Having defined the staged transition from cTregs through eTregs to exTregs, we next asked whether Treg destabilization converged on a uniform exTreg fate. Mapping the density-defined exTreg region onto the CD4+ T cell transcriptional landscape identified eight exTreg-associated clusters (Fig. 3a). Differential gene expression and CITE-seq analysis further distinguished these populations by their transcriptional and surface protein profiles (Fig. 3b), indicating substantial molecular heterogeneity within the exTreg compartment.

**Fig. 3.**
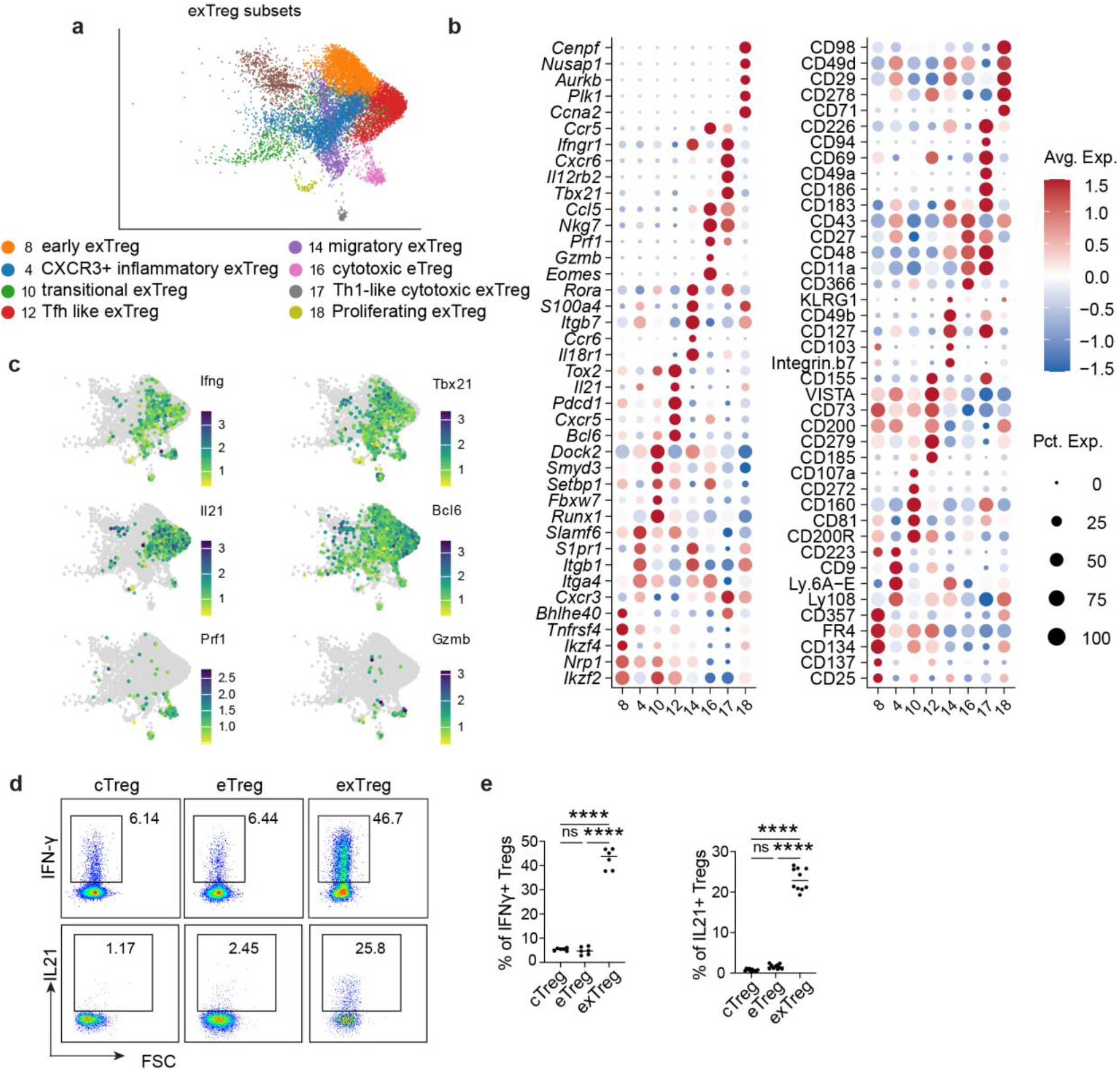
Foxp3-lineage exTreg cells comprise transcriptionally and phenotypically distinct states. **a,** UMAP visualization of exTreg-associated clusters identified from the CD4+ T cell clusters defined in Fig. 1g based on their overlap with the density-defined exTreg region shown in Fig. 2a and Extended Data Fig. 2b. Clusters are colored and numbered according to their original CD4+ T cell cluster identities. **b,** Dot plots showing five representative genes and antibody-derived tags (ADTs) selected from the top 15 differentially expressed genes and ADTs, respectively, for each exTreg-associated cluster. Dot size indicates the percentage of cells expressing each feature and color indicates scaled average expression. **c,** UMAP feature plots showing expression of *Ifng* and *Tbx21*, *Il21* and *Bcl6*, and *Prf1* and *Gzmb* in exTreg cells. **d,** Representative flow cytometry plots showing IFN-γ and IL-21 expression in cTregs, eTregs and exTregs. **e,** Frequencies of IFN-γ+ and IL-21+ cells among cTregs, eTregs and exTregs. Each symbol represents an individual mouse (*n* ≥ 6 mice per group). Data are presented as mean ± SEM. Statistical significance was determined by paired one-way ANOVA with multiple-comparison correction. \**P* < 0.05, \*\**P* < 0.01, \*\*\**P* < 0.001, **\*\*\*\****P* < 0.0001, ns, not significant.

Among these populations, cluster 8 retained activated Treg features, including *Tnfrsf4*, *Tnfrsf9*, *Ikzf2*, *Ikzf4* and *Nrp1*, while acquiring inflammatory genes such as *Cst7*, *Bhlhe40*, *Cxcl10* and *Irf5*, consistent with an early exTreg state (Extended Table. 1). Residual expression of CD25, CD137, OX40 and FR4 at the protein level further supported this identity (Fig. 3b and Extended Data Fig. 4). Cluster 10 lacked a dominant effector program and instead preferentially expressed transcriptional regulators including *Arid1b*, *Runx1*, *Setbp1* and *Fbxw7*, together with a distinctive surface phenotype marked by CD200R, CD81, CD160 and CD272, consistent with a transitional exTreg state. Cluster 4 was distinguished by expression of Cxcr3 together with activation- and migration-associated genes including *S1pr1*, *Itga4* and *Itgb1*, consistent with an Th1-like inflammatory phenotype. Expression of *Itga4* and *Itgb1*, encoding the α4β1 integrin (VLA-4), the major receptor for VCAM-1 in atherosclerosis, suggested these cells may be particularly suited for migration into inflamed vascular lesions^41, 42, 43^. Preferential expression of CD223 and Ly6A/E at the protein level further supported this CXCR3⁺ inflammatory exTreg state (Fig. 3b and Extended Data Fig. 4). These populations represent progressive stages of regulatory loss accompanied by increasing inflammatory remodeling.

Other exTreg populations displayed more specialized effector programs. Cluster 12 expressed the classical Tfh-associated genes *Cxcr5*, *Bcl6*, *Pdcd1*, *Il21* and *Tox2*, together with preferential surface expression of CXCR5 and PD-1, defining a Tfh-like exTreg state (Fig. 3b and Extended Data Fig. 4). Immediately adjacent to cluster 4, cluster 14 retained the inflammatory program while additionally expressing *Ccr6* and *Itgb7*, the β subunit of gut-homing integrins, together with CD103 and CD49b, consistent with a highly migratory phenotype. Two additional populations acquired cytotoxic programs. Cluster 16 highly expressed *Nkg7*, *Prf1*, *Gzmb* and *Ccl5.* Cluster 17 shared this cytotoxic program while additionally expressing Th1-associated genes including *Tbx21*, *Cxcr6* and *Il12rb2*, distinguishing cytotoxic and Th1-like cytotoxic exTreg states, respectively. Cluster 18 was dominated by mitotic genes including *Ccna2*, *Ccnb2*, *Plk1* and *Aurkb* together with high expression of CD71 and CD98, identifying a proliferating exTreg population.

Several of these states resembled exTreg phenotypes previously described in inflammatory and autoimmune settings. Feature mapping of representative lineage-defining genes showed that Th1-associated (*Ifng*, *Tbx21*), Tfh-associated (*Il21*, *Bcl6*) and cytotoxic (*Prf1*, *Gzmb*) programs occupied distinct yet partially overlapping regions of the exTreg landscape (Fig. 3c). Rather than forming sharply separated populations, these expression patterns suggested gradual transitions between exTreg states.

To determine whether these transitions reflected a continuous differentiation program, we modeled gene expression dynamics along pseudotime using tradeSeq^44^. Dynamic genes resolved into six sequential transcriptional modules (Extended Data Fig. 3c). Early modules corresponded to resting and proliferative and transitional Treg programs, followed by activated Treg programs enriched in eTregs, whereas late modules captured inflammatory, cytotoxic and terminal Tfh-like differentiation. Projection of these modules onto the UMAP demonstrated their sequential activation across the cTreg–eTreg–exTreg continuum (Extended Data Fig. 3d).

Flow cytometric analysis further confirmed progressive acquisition of inflammatory effector functions, with IFN-γ- and IL-21-producing cells increasing from cTregs to eTregs and being most abundant in exTregs (Fig. 3d, e). In contrast, we did not detect a prominent Th17-like exTreg population in this model (Extended Data Fig. 5). Together, these analyses indicate that Treg destabilization is a continuous differentiation process that progressively diversifies into multiple specialized effector exTreg states rather than converging on a single terminal fate.

### exTreg states emerge along distinct differentiation trajectories

The molecular heterogeneity of exTregs raised the question of whether these populations represented independent terminal states or diverging differentiation pathways. We therefore applied Slingshot trajectory inference^45^ using early exTregs (cluster 8) as the root population. Slingshot identified four branching lineages that shared an initial trajectory through transitional cluster 10 before diverging toward cytotoxic (cluster 16), Th1-like cytotoxic (cluster 17), Tfh-like (cluster 12) and proliferating (cluster 18) exTreg states (Fig. 4a). Dominant-lineage pseudotime placed cluster 8 at the earliest stage, followed sequentially by clusters 10, 4 and 14, whereas clusters 12, 16, 17 and 18 occupied the terminal regions of their respective trajectories (Fig. 4b). Together, these analyses define a branched differentiation architecture in which a shared transitional exTreg state precedes diversification into multiple terminal effector fates.

**Fig. 4.**
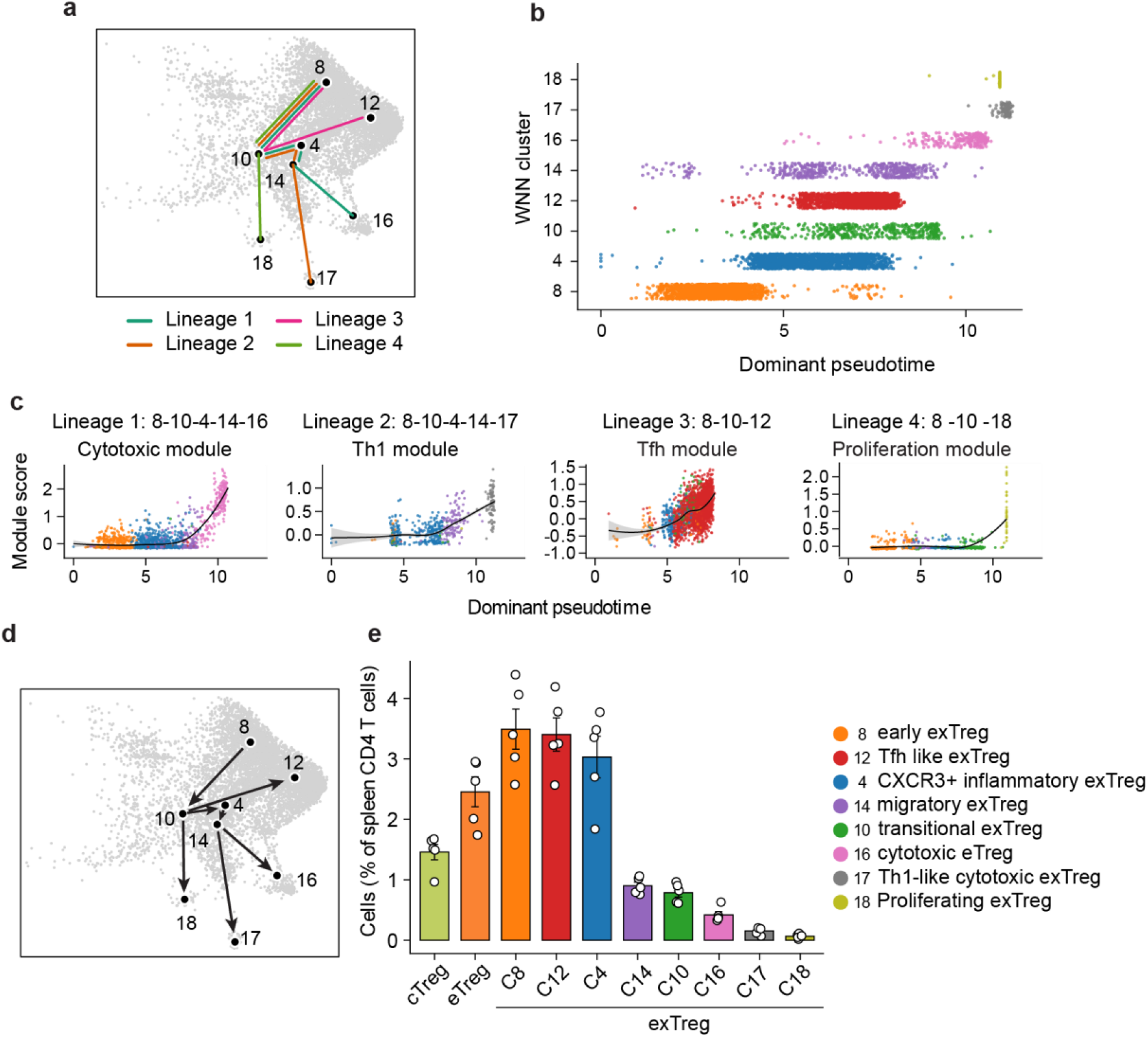
exTreg states emerge along distinct differentiation trajectories. **a,** Cluster-level topology of Slingshot-inferred exTreg lineages in WNN UMAP space. Colored lines indicate the four inferred lineages originating from cluster 8. **b,** Distribution of dominant-lineage pseudotime across WNN clusters. For each cell, the dominant lineage was assigned as the lineage with the highest Slingshot curve weight, and pseudotime was obtained from the corresponding lineage. Points represent individual cells and are colored by WNN cluster identity. **c,** Representative transcriptional programs associated with the four inferred exTreg trajectories. Cytotoxic, Th1, Tfh and proliferation module scores are shown along Slingshot pseudotime for lineages 1, 2, 3 and 4, respectively. Points represent individual cells and are colored by WNN cluster identity. Black lines indicate LOESS fits and shaded areas indicate 95% confidence intervals. **d**, Schematic summary of the exTreg differentiation trajectories inferred from the analyses in **a**–**c**. Arrows depict the shared progression from cluster 8 through transitional cluster 10, followed by divergence toward cytotoxic, Th1-like cytotoxic, Tfh-like and proliferating exTreg states. **e**, Frequency of cTregs, eTregs and individual exTreg subsets among splenic CD4 T cells. Bars represent mean ± s.e.m., and dots represent individual mice.

Lineage-specific transcriptional modules displayed distinct activation patterns along pseudotime (Fig. 4c). Cytotoxic, Th1, Tfh and proliferation programs accumulated selectively along lineages 1, 2, 3 and 4, respectively, linking each developmental branch to a distinct effector program.

Although lineages 1 and 2 shared an early differentiation path, they ultimately generated two phenotypically distinct cytotoxic exTreg populations. tradeSeq identified the molecular programs underlying this late lineage divergence (Fig. 5a). Both lineages upregulated a small set of shared cytotoxic genes, including *Ccl5*, whereas most dynamic genes were lineage specific. Lineage1 acquired a cytotoxic effector program, whereas lineage2 diverge to Th1-associated effector program. Similarly, lineage 3 and 4 share activation-, inflammatory- and migration associated genes (*Tox*, *Ifngr1*, *Hif1a*, *Cxcr3*) (Fig. 5b). Lineage3 and 4 then differentiate toward Tfh and proliferative program respectively.

**Fig. 5.**
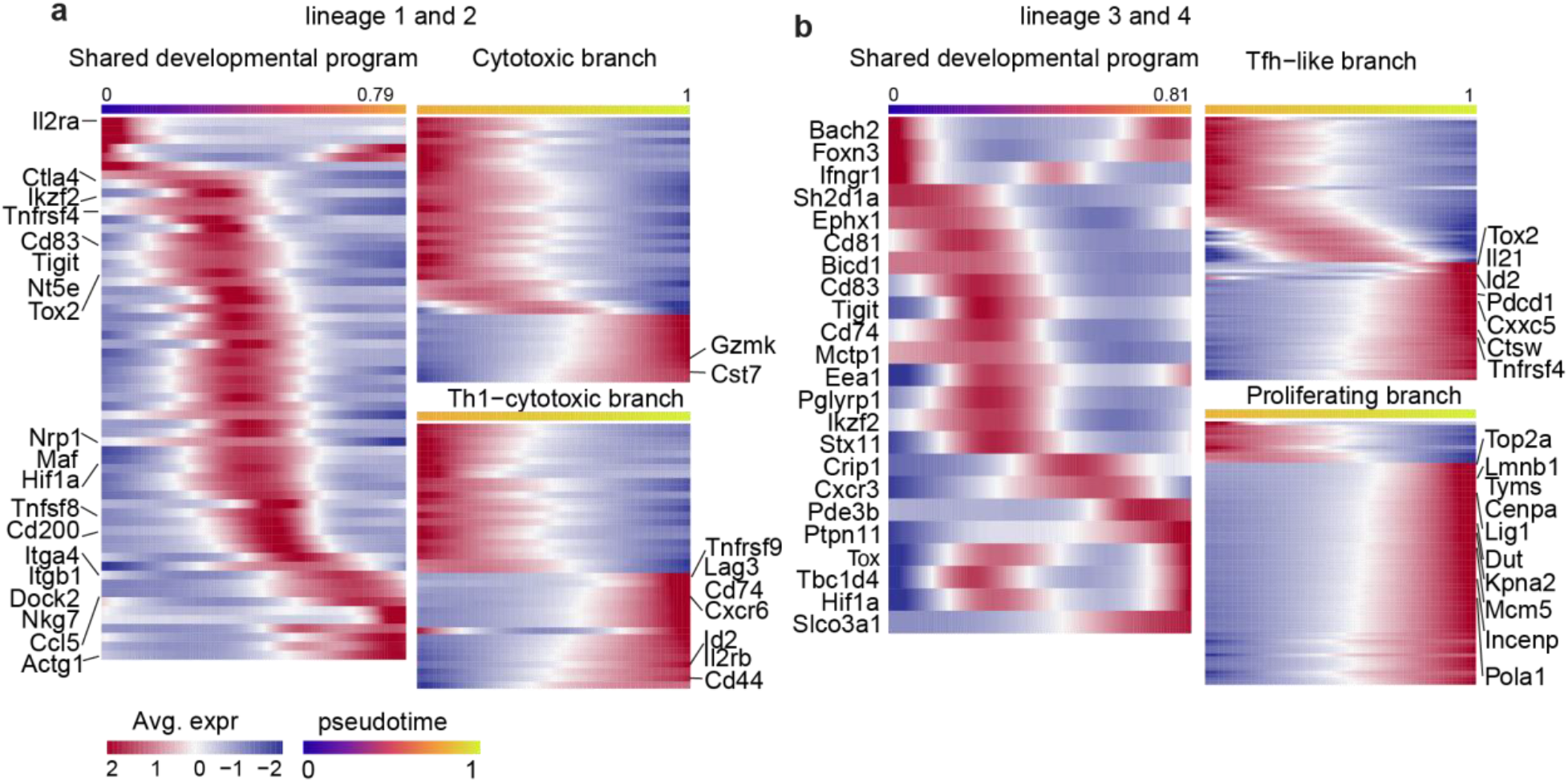
Lineage-specific gene expression dynamics along cytotoxic exTreg trajectories. **a,** Dynamic genes shared between the cytotoxic and Th1-cytotoxic trajectories (left) and genes specifically enriched in each terminal branch (right) identified by tradeSeq analysis. **b**, Dynamic genes shared between the Tfh-like and proliferating trajectories (left) and branch-enriched genes defining each terminal lineage (right). Smoothed gene expression was inferred using the joint multi-lineage tradeSeq model and displayed after row-wise Z-score normalization. Shared genes were visualized using the averaged fitted expression profiles from both lineages prior to the branch point, whereas branch-enriched genes were visualized using the post-branch portions of each lineage. Pseudotime was normalized to the complete lineage (0–1), allowing direct comparison of the relative timing of transcriptional changes across the shared and branch-specific programs.

Lineage-associated transcription factors displayed branch-specific dynamics (Extended Data Fig. 6). The cytotoxic lineage preferentially upregulated *Eomes*, *Maf* and *Ikzf3*, whereas the Th1-like cytotoxic lineage progressively induced *Tbx21*, *Satb1* and *Id2*. Likewise, the Tfh lineage showed increasing expression of *Bcl6*, *Tox2* and *Pou2af1*, while the proliferative lineage displayed distinct dynamics of *Ybx3*, *Ikzf3* and *Foxp1*. These transcription factors represent candidate regulators of exTreg lineage diversification.

Together, these findings establish a branched differentiation model in which a shared transitional exTreg state gives rise to multiple specialized effector fates through lineage-specific transcriptional programs.

### Tissue context shapes exTreg accumulation and composition

Because the axillary and cervical lymph nodes drain the tissues surrounding the diseased aorta and carotid arteries, we asked whether exTreg accumulation differed between draining and non-draining lymph nodes. cTregs and eTregs distributions were similar between the two lymph node compartments, whereas exTregs preferentially accumulated in dLNs, with both cell number and frequency increasing from ndLNs to dLNs (Fig. 6a, b). Thus, the staged Treg-lineage states identified in the spleen were conserved across lymphoid tissues, whereas exTreg accumulation varied substantially with tissue location.

**Fig. 6.**
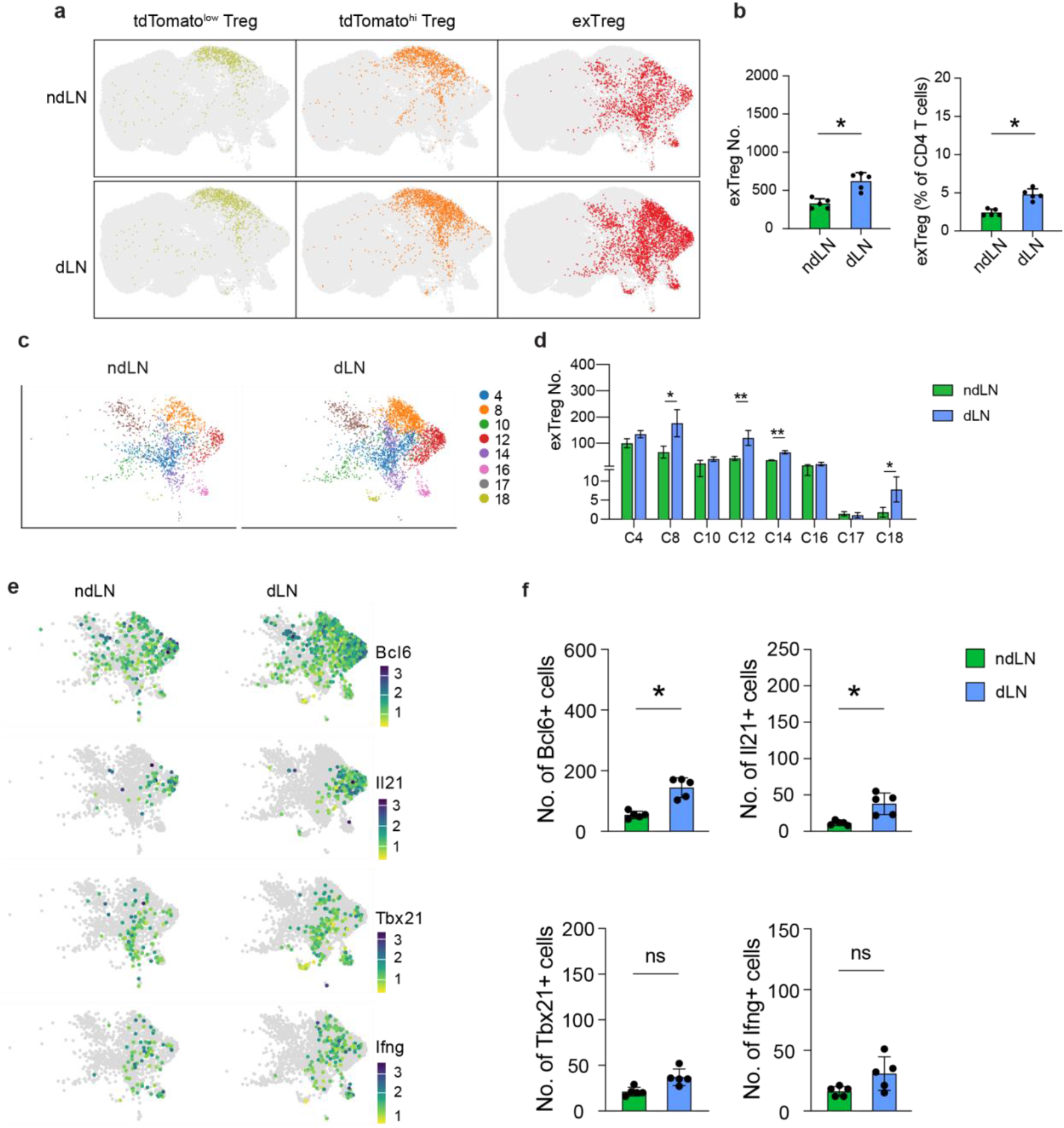
exTreg states differentially accumulate across lymphoid tissues. **a,** WNN UMAPs showing the distribution of tdTomato^low^ Tregs (cTregs), tdTomato^hi^ Tregs (eTregs) and exTregs among CD4+ T cells from non-draining lymph nodes (ndLNs) and draining lymph nodes (dLNs). Cells belonging to the indicated population are colored and all other CD4+ T cells are shown in gray. **b,** Number (left) and frequency among CD4+ T cells (right) of exTregs in ndLNs and dLNs. Each point represents one mouse. **c,** WNN UMAPs showing the distribution of exTreg-associated clusters in ndLNs and dLNs. Cells are colored by WNN cluster identity. **d,** Number of exTreg cells in each WNN cluster in ndLNs and dLNs. Each bar represents the mean across mice and error bars indicate s.e.m. **e,** WNN UMAP feature plots showing expression of *Bcl6*, *Il21*, *Tbx21*, and *Ifng* in exTregs from ndLNs and dLNs. **f,** Numbers of *Bcl6*+, *Il21*+, *Tbx21*+, and *Ifng*+ exTregs in ndLNs and dLNs. Each point represents one mouse. Data in **b,f** are presented as mean ± SEM. n = 5 mice. Statistical significance was determined by paired one-way ANOVA followed by Tukey’s multiple-comparison test. \**P* < 0.05, \*\**P* < 0.01, \*\*\**P* < 0.001, **\*\*\*\****P* < 0.0001, ns, not significant.

We next examined whether increased exTreg accumulation was accompanied by changes in exTreg composition. All major exTreg states identified in the spleen were also present in ndLNs and dLNs (Fig. 6c). Their abundance generally increased from ndLNs to dLNs, particularly for early (cluster 8) and Tfh-like (cluster 12) exTregs (Fig. 6d). Relative composition further differed between lymph node compartments, with early, Tfh-like and proliferating exTregs enriched in dLNs, whereas CXCR3⁺ inflammatory exTregs were proportionally more abundant in ndLNs (Extended Data Fig. 7). This distribution is consistent with the elevated expression of CXCR3 and α4β1 integrin in this subset (Fig. 3b), suggesting enhanced migratory capacity.

Consistent with these differences, *Bcl6*- and *Il21*-expressing exTregs were enriched in dLNs, whereas *Tbx21*- and *Ifng*-expressing exTregs showed a similar but less pronounced pattern (Fig. 6e, f). Together, these findings indicate that tissue-specific exTreg accumulation is accompanied by selective enrichment of distinct effector programs, particularly the Tfh-like exTregs.

### Clonal relationships support staged exTreg differentiation and diversification

Having established staged Treg-to-exTreg differentiation and subsequent exTreg diversification, we next asked whether TCR clonotype relationships provided independent support for this developmental model. Shared clonotypes were largely restricted to tissues from the same mouse, with minimal sharing between animals (Extended Data Fig. 8a). Because clonal expansion reflects antigen-driven proliferation, we first examined its distribution across lymphoid tissues. Consistent with the tissue-specific accumulation of exTregs, expanded clonotypes of all size classes were enriched in draining compared with non-draining lymph nodes and were most abundant in the spleen (Extended Data Fig. 8b, c). Across lineage-traced Treg populations, expanded clonotypes increased progressively from cTregs to eTregs and were most abundant in exTregs (Fig. 7a, b). Representative shared clonotypes expanded progressively from cTregs to eTregs and further into exTregs (Fig. 7c). These findings provide independent clonal support for a staged cTreg-to-eTreg-to-exTreg transition.

**Fig. 7.**
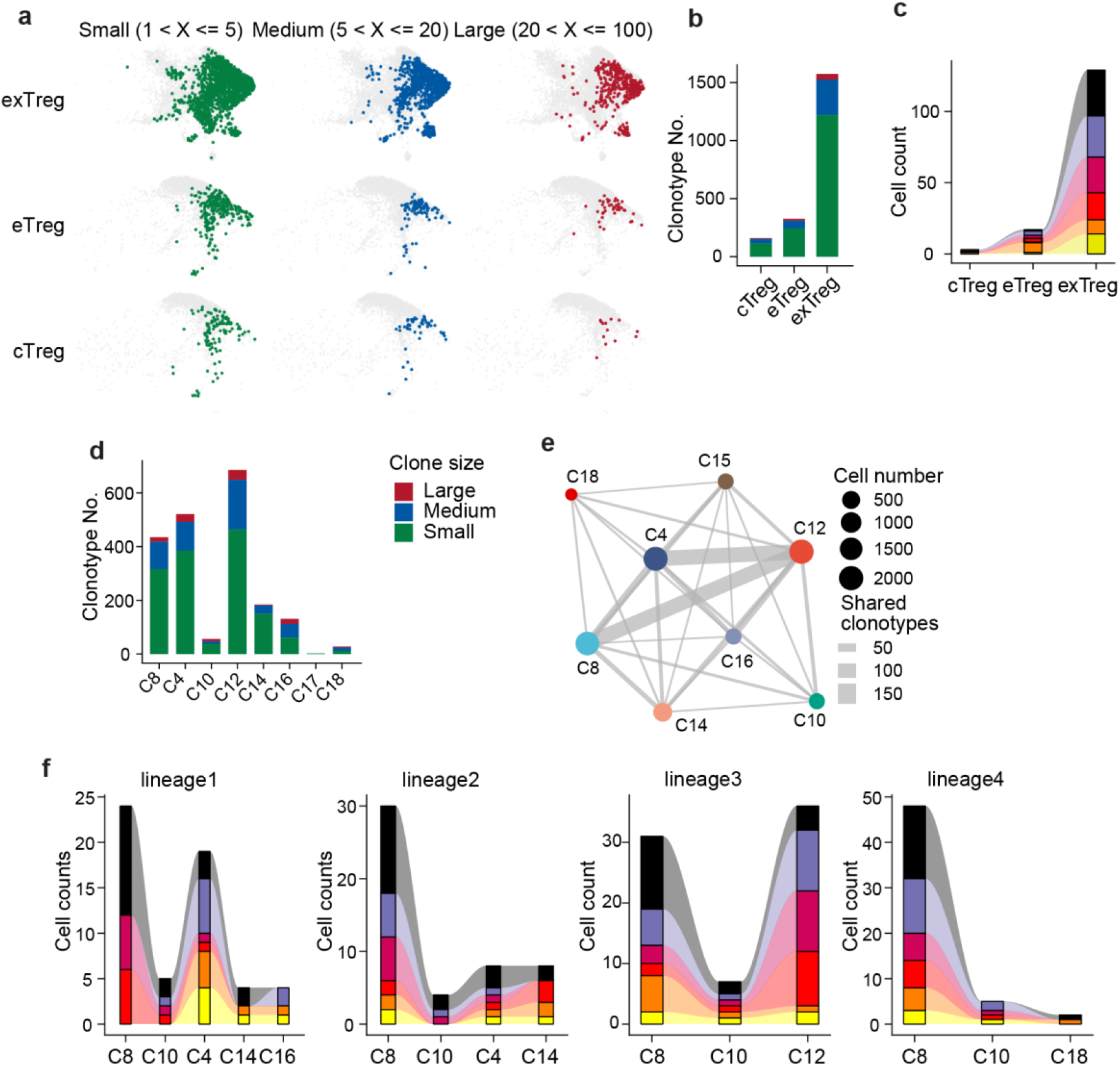
Clonal relationships support staged exTreg differentiation and diversification. **a,** WNN UMAPs showing the distribution of clonally expanded cTregs, eTregs and exTregs across the combined ndLN, dLN and spleen datasets. Clonotypes were classified according to clone size as small (1 < x ≤ 5), medium (5 < x ≤ 20) or large (20 < x ≤ 100). Cells belonging to the indicated clone-size category are colored and other cells are shown in gray. **b,** Number of small, medium and large expanded clonotypes in cTregs, eTregs and exTregs. **c,** Alluvial plot showing the six most abundant clonotypes shared across cTregs, eTregs and exTregs. Colored flows represent individual shared clonotypes, and flow width is proportional to the number of cells carrying each clonotype within the indicated Treg state. **d,** Number of small, medium and large expanded clonotypes across exTreg clusters. **e,** Clonal sharing network among exTreg clusters. Nodes represent exTreg clusters, with node size proportional to cell number. Edges connect clusters containing shared clonotypes, and edge width is proportional to the number of shared clonotypes. **f,** Distribution of representative shared clonotypes across exTreg clusters along each of the four Slingshot-inferred lineages. Stacked areas and bars indicate the numbers of cells carrying individual shared clonotypes in each cluster. Clusters are ordered according to the corresponding Slingshot-inferred lineage: lineage 1, 8-10-4-14-16; lineage 2, 8-10-4-4-17; lineage 3, 8-10-12; and lineage 4, 8-10-18. Clonotypes were defined using the scRepertoire strict clonotype definition, requiring identical paired TCRα and TCRβ CDR3 amino acid sequences and matching V and J gene usage.

Within the exTreg compartment, clonal expansion was unevenly distributed across individual exTreg states. Expanded clonotypes were most abundant in early (C8), CXCR3⁺ inflammatory (C4) and Tfh-like (C12) exTregs, with the Tfh-like population exhibiting the greatest overall clonal representation (Fig. 7d). Large expanded clones were particularly enriched in the Tfh-like and cytotoxic exTregs. In contrast, transitional, migratory, cytotoxic, Th1-like cytotoxic and proliferating exTreg states contained comparatively fewer expanded clonotypes. Shared-clonotype network analysis revealed extensive overlap centered on early, CXCR3⁺ inflammatory and Tfh-like exTregs, with the Tfh-like population acting as a major clonotypic hub (Fig. 7e; Extended Data Fig. 8d). Finally, mapping shared clonotypes onto the Slingshot-inferred differentiation paths demonstrated that individual clonotypes were distributed across successive clusters within each lineage rather than being confined to terminal states (Fig. 7f). Together, these analyses provide independent clonal evidence supporting both the staged cTreg-to-eTreg-to-exTreg transition and the subsequent diversification of exTregs into distinct effector states.

### IL-6–IL-6R signaling promotes Treg-to-exTreg conversion during atherosclerosis

The preceding analyses established Treg destabilization as a staged and branched differentiation process, but the inflammatory signals promoting this transition remained unclear. IL-6–IL-6R signaling has been implicated in impaired Treg stability through STAT3-dependent disruption of Foxp3 maintenance^34, 46^. We therefore examined whether this pathway promoted Treg-to-exTreg conversion during atherosclerosis.

Plasma IL-6 was elevated in Western diet-fed *Apoe*^−/−^ mice (Fig. 8a). *Il6ra* was broadly expressed throughout the Foxp3-lineage compartment, with enrichment in exTregs confirmed by both single-cell transcriptomics and CD126 flow cytometry (Fig. 8b–d). We therefore asked whether IL-6 directly promotes Treg-to-exTreg conversion in vitro. In vehicle-treated cultures, most cells differentiated into Foxp3⁺tdTomato^hi^ eTregs, whereas a smaller fraction became GFP⁻tdTomato⁺ exTregs by day 6 (Fig. 8e). Addition of IL-6 significantly increased exTreg generation, whereas IL-6 neutralization restored exTreg frequencies to control levels. This difference became even more pronounced by day 10. Notably, exTregs were still generated in vehicle-treated cultures and after IL-6 neutralization, indicating that Treg-to-exTreg conversion can also occur through IL-6-independent mechanisms.

**Fig. 8.**
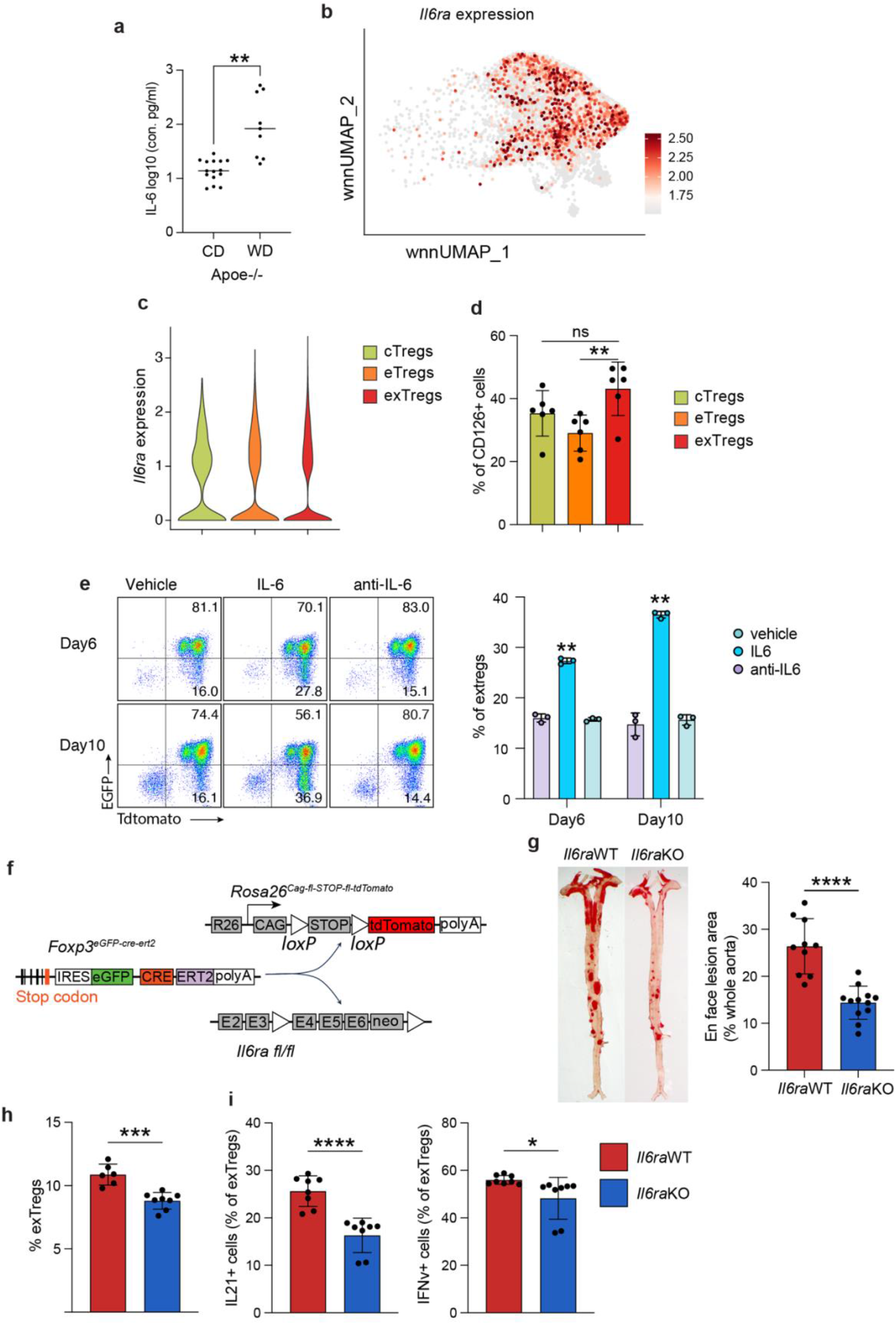
Treg-intrinsic IL-6R signaling promotes Treg-to-exTreg conversion and atherosclerosis. **a,** Plasma IL-6 concentrations in *Apoe*^−/−^ mice fed a chow diet (CD) or western diet (WD). Plasma was collected after 20 weeks of diet feeding and IL-6 concentrations were measured. **b,** WNN UMAP feature plot showing *Il6ra* expression across combined cTreg, eTreg and exTreg cells from ndLN, dLN and spleen. **c,** Violin plots showing *Il6ra* expression in cTregs, eTregs and exTregs. **d,** Flow cytometric quantification of surface IL-6Rα (CD126) expression in splenic cTregs, eTregs and exTregs from Foxp3 lineage-tracing *Apoe*^−/−^ mice after 20 weeks of western diet feeding. the percentage of CD126+ cells is shown. **e,** In vitro conversion of current Tregs into exTregs in response to IL-6. FACS-sorted Foxp3+tdTomato+ current Tregs, comprising cTregs and eTregs, were cultured with IL-2 (500 U/ml) and anti-CD3/CD28 beads in the presence of vehicle, IL-6 (50 ng/ml) or an IL-6-neutralizing antibody (anti-IL-6; 50 ng/ml) for 10 d. Representative flow cytometry plots showing Foxp3-EGFP and tdTomato expression at days 6 and 10 (left) and quantification of Foxp3−tdTomato+ exTregs (right). *n* = 3 independent cultures per condition. **f,** Schematic of the breeding strategy used to generate inducible Treg-specific *Il6ra*-deficient Foxp3 lineage-tracing mice. *Foxp3*^eGFP-Cre-ERT2^ *Rosa26*^CAG-flox-STOP-flox-tdTomato^ mice were crossed with *Il6ra* fl/fl mice. **g,** Representative Sudan IV-stained whole aortas from Treg-specific *Il6ra* wild-type (*Il6ra*WT) and *Il6ra*-deficient (*Il6ra*KO) Foxp3 lineage-tracing *Apoe*^−/−^ mice after tamoxifen administration and 20 weeks of western diet feeding (left), and quantification of *en face* atherosclerotic lesion area as a percentage of total aortic surface area (right). **h,** Frequency of splenic exTregs among CD4+ T cells in *Il6ra*WT and *Il6ra*KO Foxp3 lineage-tracing *Apoe*^−/−^ mice after 20 weeks of western diet feeding. **i,** Frequencies of IL-21+ (left) and IFN-γ+ (right) cells among splenic exTregs from *Il6ra*WT and *Il6ra*KO mice. Splenic CD4+ T cells were stimulated with PMA and ionomycin for 6 h; protein transport inhibitor was added after 2 h, and cytokine production was assessed by intracellular flow cytometry. Each symbol represents an individual mouse unless otherwise indicated. Data are presented as mean ± SEM. Statistical significance was determined by one-way ANOVA followed by Tukey’s multiple-comparison test or by unpaired two-tailed Student’s *t*-test. \**P* < 0.05, \*\**P* < 0.01, \*\*\**P* < 0.001, **\*\*\*\****P* < 0.0001, ns, not significant.

To determine whether Treg-intrinsic IL-6R signaling contributed to Treg destabilization in vivo, we induced Treg-specific deletion of *Il6ra* in Foxp3 lineage-tracing *Apoe*^−/−^ mice (Fig. 8f). Il6ra deficiency reduced atherosclerotic lesion size (Fig. 8g). This was accompanied by reduced frequencies of splenic exTregs, particularly IL-21- and IFNγ-expressing exTregs (Fig. 8h, i). Together, these findings identify Treg-intrinsic IL-6R signaling as a key regulator of Treg-to-exTreg conversion and inflammatory exTreg differentiation during atherosclerosis.

## Discussion

Treg instability is increasingly recognized in chronic inflammatory and autoimmune diseases^26, 27, 28^, yet how regulatory identity is progressively lost and gives rise to diverse exTreg states remains poorly understood. Here, we define Treg destabilization during atherosclerosis as a staged and branching differentiation process rather than an abrupt loss of lineage identity. Conventional Tregs first transition through an effector Treg state before entering a heterogeneous exTreg compartment comprising eight transcriptionally distinct subsets, including Tfh-like, cytotoxic, Th1–cytotoxic and proliferative exTregs. Transcriptional trajectories, TCR clonotype relationships and experimental Treg-to-exTreg conversion converged on this developmental framework. We further identify Treg-intrinsic IL-6R signaling as a promoter of exTreg differentiation and link reduced exTreg accumulation to attenuated atherosclerosis. Together, these findings shift the view of Treg instability from a binary loss of identity toward a staged differentiation process through which chronic inflammation redirects regulatory T cell fate.

The identification of eTregs as an intermediate state has important implications for how Treg plasticity and instability are viewed. Effector differentiation is an integral component of physiological Treg adaptation^18, 21, 22, 24, 47^, enabling Tregs to acquire specialized migratory and functional properties while maintaining regulatory identity. By contrast, Treg instability is typically defined by loss of Foxp3 and acquisition of inflammatory effector function^26, 27, 28^. Our findings suggest that these two processes may be developmentally connected during chronic vascular inflammation. Compared with cTregs, eTregs exhibited progressive attenuation of the CD25–STAT5 axis and Treg-associated transcriptional programs while acquiring activated and antigen-experienced features, placing them between cTregs and exTregs. This intermediate phenotype, together with trajectory inference and experimental cTreg-to-eTreg-to-exTreg conversion, supports a model in which Treg identity is progressively remodeled rather than being lost through single discrete transition. Importantly, these findings do not imply that eTregs are obligatorily committed to an exTreg fate. Stable eTregs are essential components of immune regulation ^22, 24^, and the factors that determine whether eTregs maintain regulatory function or progress toward destabilization remain incompletely understood.

A second major implication of our findings is that loss of Foxp3 does not result in a uniform exTreg state. Instead, our data indicate that the exTreg compartment comprises multiple specialized effector states organized along distinct differentiation trajectories. Previous studies identified IFN-γ-producing Th1-like Treg-lineage cells and Treg-to-Tfh conversion in atherosclerosis, whereas cytotoxic exTregs were identified in both mouse and human cardiovascular disease^9, 10, 11, 12^. Rather than representing independent examples of Treg plasticity reported in isolation, our findings suggest that multiple previously described exTreg phenotypes can arise through a shared process of exTreg diversification. This diversification was accompanied by progressively distinct effector programs and lineage-associated transcriptional regulators, suggesting continued specialization after entry into the exTreg compartment. In addition, our analyses identify early, transitional, migratory and proliferative exTreg states that have not previously been recognized. The limited representation of Th17-like exTregs in our atherosclerosis model, despite their prominence in other inflammatory settings, further suggests that the outcome of exTreg differentiation is shaped by disease-specific inflammatory cues. Thus, exTregs may be better viewed as a heterogeneous compartment of specialized effector states rather than a single terminal outcome of Treg destabilization.

TCR clonotype analysis provided independent evidence supporting the proposed model of exTreg differentiation. Shared clonotypes spanning cTregs, eTregs and multiple exTreg states supported developmental continuity across the inferred differentiation process, whereas clonotype sharing among transcriptionally distinct exTreg subsets suggested that divergent effector programs can emerge from clonally related Treg-lineage cells. Nevertheless, clonal relationships did not always mirror transcriptional trajectories. For example, extensive clonotype sharing between CXCR3+ inflammatory and Tfh-like exTregs indicates that clonal expansion and transcriptional specialization are not necessarily coupled processes. The predominantly private nature of expanded clonotypes further suggests a polyclonal antigen response rather than expansion driven by a single dominant specificity. Together, these findings support a model in which Treg destabilization generates a clonally connected but developmentally flexible exTreg compartment, allowing related Treg-lineage cells to adopt distinct effector programs.

The tissue distribution of exTregs further suggests that immune context may influence their accumulation and differentiation. ExTregs accumulated progressively from non-draining lymph nodes to draining lymph nodes and spleen, accompanied by increasing clonal expansion across these compartments. This distribution is consistent with disease-associated immune environments that favor exTreg accumulation and possibly Treg destabilization in both vascular-draining and systemic lymphoid tissues. Notably, Tfh-like exTregs represented the dominant differentiated exTreg state and also accumulated across these tissue compartments. ExTregs were also more clonally expanded than cTregs or eTregs, raising the possibility that antigen-associated clonal expansion contributes to their emergence or accumulation. Although antigen specificity was not resolved in our dataset, previous studies have shown that ApoB-reactive Tregs undergo clonal expansion together with progressive remodeling of Treg identity during atherosclerosis, including acquisition of inflammatory transcriptional programs, gradual Treg-to-exTreg conversion and loss of the effector Treg phenotype^4, 5, 33, 48^. However, whether ApoB or other self-antigens contribute to the expansion or diversification of exTreg clones remains unknown. Together, the tissue distribution and clonal structure of exTregs suggest that local immune environments shape both the extent of Treg destabilization and the composition of the resulting exTreg compartment. The mechanisms underlying this tissue-associated differentiation have yet to be defined.

Building on previous studies linking IL-6 signaling to impaired Treg stability^27, 32, 34, 46, 49, 50^, our findings place Treg-intrinsic IL-6R signaling within a broader developmental framework encompassing multiple exTreg states. IL-6R expression increased progressively from cTregs to eTregs and exTregs, suggesting that responsiveness to IL-6 increases during Treg destabilization. Consistent with this framework, exogenous IL-6 accelerated exTreg generation in vitro, whereas Treg-specific deletion of Il6ra reduced exTreg accumulation, diminished IL-21- and IFN-γ-producing exTreg populations, and attenuated atherosclerosis in vivo. Importantly, exTregs were not completely eliminated following Treg-specific deletion of Il6ra, indicating that IL-6R signaling is an important, but not exclusive, driver of Treg destabilization. Together, the progressive loss of CD25–STAT5 signaling and increased IL-6R expression raise the possibility of a shift from lineage-stabilizing to inflammatory cytokine responsiveness during Treg destabilization. Whether this reciprocal shift directly contributes to progression through the staged exTreg differentiation program remains to be determined.

Several questions remain unresolved. Although the Foxp3 lineage-tracing system enables tracking of Treg-lineage cells, transient Foxp3 expression may label a small fraction of cells that do not represent fully committed Tregs^51, 52^. In addition, because our analyses were performed at a single stage of disease progression, the temporal evolution of exTreg differentiation remains unresolved. Therefore, additional exTreg states or alternative differentiation trajectories may emerge at earlier or later stages of atherosclerosis. Transcriptional trajectories, TCR clonotype relationships and experimental conversion support a staged model of Treg destabilization, but definitive lineage relationships will require prospective lineage tracing of defined eTreg and exTreg states. Such studies will also be needed to determine whether individual exTreg states are stable, reversible or terminal. It also remains unclear how antigen specificity influences exTreg fate or whether the same differentiation architecture is conserved in human atherosclerosis.

In summary, our study defines Treg destabilization during atherosclerosis as a staged and branching differentiation process in which conventional Tregs progress through an effector intermediate before diversifying into multiple pathogenic exTreg states. By identifying Treg-intrinsic IL-6R signaling as an important driver of this process, our findings establish a developmental framework for understanding how chronic inflammation redirects Treg cell fate and suggest that preserving Treg lineage stability may represent a therapeutic strategy in atherosclerosis and other chronic inflammatory diseases.

## Methods

### Mice

Apoe−/−, Foxp3-EGFP-Cre-ERT2 and Rosa26-CAG-LSL-tdTomato mice were originally obtained from The Jackson Laboratory (Bar Harbor, ME, USA; stock numbers. 002052, 016961 and 007909, respectively). *Il6ra*^fl/fl^ mice were kindly provided by David Munn (Augusta University, GA, USA). Foxp3-EGFP-Cre-ERT2 and Rosa26-CAG-LSL-tdTomato mice were crossed with *Apoe*^−/−^ mice to generate Foxp3 lineage-tracing Apoe^−/−^ mice. *Il6ra*^fl/fl^ mice were further crossed with Foxp3 lineage-tracing *Apoe*^−/−^ mice to generate *Il6ra*^fl/fl^ *Foxp3-EGFP-Cre-ERT2 Rosa26-CAG-LSL-tdTomato Apoe*^−/−^ mice. *Il6ra* wild-type Foxp3 lineage-tracing *Apoe*^−/−^ mice were used as controls for experiments examining Treg-intrinsic IL-6R signaling.

For Foxp3 lineage tracing, 7–8-week-old mice received tamoxifen (75 mg/kg body weight; Millipore Sigma, T2859) by intraperitoneal injection once daily for 5 consecutive days. Tamoxifen was dissolved in olive oil (Sigma-Aldrich, #O1514). Mice were then placed on a Western diet (envigo, cat#:TD.88137) and received a second 5-day course of tamoxifen after 6 weeks of Western diet feeding. Chow-fed controls received a standard chow diet (PicoLab, cat#: 5053). Mice were analyzed after 12 weeks of diet feeding.

For experiments examining Treg-intrinsic IL-6R signaling, *Il6ra*^fl/fl^ and *Il6ra* wild-type Foxp3 lineage-tracing *Apoe*^−/−^ mice received the same initial tamoxifen regimen at 7-8 weeks of age before western diet feeding. A second 5-day course of tamoxifen was administered after 10 weeks of Western diet feeding, and mice were analyzed after 20 weeks of western diet feeding.

Mice were bred and maintained in microisolator cages under specific pathogen-free conditions in an AAALAC-accredited and OLAW-assured animal facility at Augusta University. Animals were maintained on a 12-h light–dark cycle (lights on at 06:00 and off at 18:00), at an ambient temperature of 20–24 °C and relative humidity of 30–70%. All animal procedures were approved by the Augusta University Institutional Animal Care and Use Committee and were performed in accordance with the NIH Guide for the Care and Use of Laboratory Animals. Both male and female mice were used for wet-laboratory experiments and were randomly assigned to experimental groups. Female mice were used for single-cell sequencing. Investigators were blinded to experimental group during lesion quantification and, where applicable, data analysis.

### Cell isolation

Spleens, draining lymph nodes (cervical, brachial and axillary) and non-draing lymph nodes (inguinal, popliteal) were collected in staining buffer consisting of PBS supplemented with 2% fetal bovine serum (FBS; GeminiBio, S11150). Tissues were mechanically dissociated through 70-µm cell strainers and washed with staining buffer. Splenic red blood cells were lysed with 1× eBioscience RBC Lysis Buffer (ThermoFisher Scientific, 00-4333-57) for 5 min. After lysis, cells were washed with PBS, passed through 40-µm filters and counted.

Single-cell suspensions from spleens and lymph nodes were either used directly for flow cytometry or subjected to CD4+ T cell enrichment for downstream analyses. CD4+ T cells were enriched using the EasySep Mouse CD4+ T Cell Isolation Kit (STEMCELL Technologies) according to the manufacturer’s instructions. Enriched CD4+ T cell preparations were >90% pure, as determined by surface expression of TCRβ and CD4 by flow cytometry.

### Flow cytometry

Single-cell suspensions from spleens or lymph nodes, or enriched CD4 T cells were stained with a fixable viability dye (LIVE/DEAD Fixable Blue Dead Cell Stain, ThermoFisher scientific) for 30min at 4 °C, followed by Fc receptor blockade with TruStain FcX™ anti-mouse CD16/32 Antibody (BioLegend). Unless otherwise indicated, all staining procedures were performed on ice. Samples were acquired on a 5-Laser Cytek Aurora Spectral flow Cytometer (Cytek Biosciences). All flow cytometry data were analyzed using FlowJo version 10.10.0.

For surface staining, cells were stained in cold staining buffer with antibodies against CD4 (clone Gk1.5; BD Biosciences), TCRβ (clone H57-597; BioLegend), CD25 (clone PC61.5; eBiosciene), CD62L (clone MEL-14; BioLegend), CD44 (clone IM7; BioLegend), CD126 (clone D7715A7; BD Biosciences), and CD45 (clone 30-F11; BioLegend). EGFP and tdTomato fluorescence were detected directly without antibody staining unless otherwise indicated.

For phospho-STAT5 (pSTAT5) analysis, freshly isolated cells or enriched CD4+ T cells were seeded in 96-well plates and surface stained for TCRβ, CD4 and CD25 at 4 °C. Cells were then stimulated with recombinant mouse IL-2 (500 U/mL; Miltenyi biotec) in 100 µl culture medium for 20 min at 37 °C. Immediately after stimulation, 100 µl pre-warmed BD Cytofix fixation buffer (BD Biosciences) was added directly to each well, and cells were fixed for 15 min at 37 °C. Cells were centrifuged at 300 × g for 5–10 min and permeabilized with pre-chilled BD Phosflow Perm Buffer III (BD Biosciences) for 30 min at room temperature. After two washes with staining buffer, cells were stained with antibodies against pSTAT5 (clone 47/Stat5(pY694); BD Biosciences) and Foxp3 (clone FJK-16S; eBioscience) before flow cytometric analysis.

For intracellular cytokine staining, enriched CD4+ T cells were stimulated in RPMI medium supplemented with 10% FBS with 1× eBioscience Cell Stimulation Cocktail (500×; ThermoFisher Scientific) containing PMA and ionomycin for 6 h in the presence of 1× eBioscience Protein Transport Inhibitor Cocktail (500×; ThermoFisher scientific) containing Brefeldin A and Monensin. Cells were subsequently surface stained for CD4, TCRβ, CD25 as described above. Cells were fixed for 20 min with IC Fixation buffer (ThermoFisher scientific) and then permeabilized with Foxp3/Transcription Factor Staining Buffer Set (ThermoFisher scientific). Intraculullar staining was performed in permeabilization buffer at room temperature using antibodies against IFN-γ (clone XMG1.2; BioLegend), IL-21 (clone FFA21; eBioscience), and Foxp3, followed by flow cytometric analysis.

### Cell Culture

Following CD4+ T cell enrichment with the EasySep Mouse CD4+ T Cell Isolation Kit (STEMCELL Technologies), Treg populations were sorted using a Bigfoot Spectral Cell Sorter (ThermoFisher Scientific) based on EGFP and tdTomato fluorescence. For experiments comparing Treg subsets, Foxp3-EGFP+tdTomato^low^ (cTregs) and Foxp3-EGFP+tdTomato^hi^ (eTregs) were sorted separately. For experiments using the total current Treg population, Foxp3-EGFP+ cells comprising both tdTomato^low^ and tdTomato^hi^ Tregs were sorted together. Sorted Treg cells were cultured in RPMI1640 medium supplemented with 10% FBS, 2mM L-glutamine, 1mM sodium pyruvate, 10mM HEPES, 50 µM 2-mercaptoethanol, 100 U/ml penicillin, 100 µg/ml streptomycin as indicated. Cells were activated with anti-CD3/CD28 antibody-coated beads (Life Technologies) and cultured with recombinant mouse IL-2 (200 U/mL, Miltenyi Biotec) at 37°C, 5% CO_2_. For cell division tracking, sorted Treg cells were labeled in vitro with CellTrace Violet (Life Technologies) according to the manufacturer’s instructions before culture. Cell proliferation was assessed after 6 days by CellTrace Violet dilution using flow cytometry. For Treg stability assays, recombinant mouse IL-6 (50ng/ml) was added to the cultures as indicated. Foxp3 expression and exTreg frequency were assessed by flow cytometry on days 6 and 10.

### Quantification of atherosclerosis

The entire aorta, including the thoracic and abdominal regions, was excised, cleaned in situ and fixed in 4% paraformaldehyde at 4 °C for at least 24 h. Aortas were opened longitudinally, pinned en face and stained with Sudan IV. Atherosclerotic lesion burden was quantified as the percentage of Sudan IV-positive area relative to the total surface area of the aortic arch or entire aorta. Image analysis was performed using QuPath (v.0.5.1) ^53^ by an investigator blinded to experimental group.

### Single-cell RNA-sequencing

Spleens, draining lymph nodes and non-draining lymph nodes were collected from five female Foxp3 lineage-tracing Apoe−/− mice after 12 weeks of Western diet feeding, yielding 15 biological samples. Enriched CD4+ T cells from there samples were Fc blocked and stained with the TotalSeq-C Mouse Universa Cocktail v1.0 (BioLegend) according to the manufacturer’s instructions. Cell viability was >90%. After washing, cell suspensions were adjusted to a concentration of 700-1200 cells/µl. Cell suspensions were loaded onto Chromium Next GEM Chips according to the manufacturer’s protocol. Gene expression, antibody-derived tag and T cell receptor libraries were generated using the Chromium Single Cell platform with Feature Barcode technology (10x Genomics) according to the manufacturer’s protocols. Library quality was assessed using a TapeStation system, and libraries were sequenced on an Illumina NovaSeq platform.

### Quality control after sequencing and data processing

Cell Ranger (10x Genomics) was used for demultiplexing, which yielded 11,540–19,715 cells per sample (median, 14,229 cells), comprising a total of 222,655 cells with transcriptomes across 15 samples. Gene expression reads were aligned to a custom mouse reference genome containing the tdTomato transgene sequence. Gene expression and antibody-derived tag (ADT) count matrices were imported into Seurat v5, and matched by shared cell barcodes for each sample. Cells with 250–6,000 detected genes, 500–25,000 RNA unique molecular identifier (UMI) counts and ≤5% mitochondrial transcripts were retained. ADT quality control required ≥30 detected antibody features and 500–80,000 ADT UMI counts per cell. Doublets were identified independently in each sample using scDblFinder and excluded from subsequent analyses.

TCR annotations were obtained from Cell Ranger V(D)J outputs and matched to individual cells by barcode. Only high-confidence, productive TRA and TRB contigs assigned to cells were retained. For cells with multiple contigs of the same TCR chain, the contig with the highest UMI count was selected, with read count used to resolve ties. TCR CDR3 amino acid and nucleotide sequences, V(D)J gene usage and clonotype identities were incorporated into the corresponding Seurat metadata for downstream clonal analyses.

RNA and ADT data were integrated across samples using Harmony and analyzed using the weighted nearest neighbor (WNN) framework in Seurat. ADT expression thresholds were determined individually for each antibody using ThresholdR and reviewed based on marker expression distributions^36^. CD4+ T cells were identified based on thresholded CD3 and CD4 expression and retained for downstream analyses.

CD4+ T cells were reanalyzed using Harmony integration and WNN based on the first 15 RNA and 10 ADT dimensions, followed by UMAP visualization and Leiden clustering at a resolution of 0.7. To define Treg lineage populations, normalized tdTomato RNA expression was modeled using ThresholdR. The tdTomato expression distribution was fitted using mixture models, with the number of components selected on the basis of the Bayesian information criterion. The resulting bimodal distribution was used to define tdTomato^low^ (normalized expression, 0.74–1.35) and tdTomato^hi^ (≥1.35) populations; tdTomato expression below 0.74 was considered background and excluded from tdTomato-positive populations. Foxp3 and tdTomato expression were jointly used to classify Foxp3+tdTomato^low^ Tregs (cTregs), Foxp3+tdTomato^hi^ Tregs (eTregs) and Foxp3−tdTomato+ exTregs for downstream analyses.

Cluster-specific RNA and ADT markers were identified using the Seurat FindAllMarkers function with the Wilcoxon rank-sum test and filtered for adjusted P value <0.05, average log2 fold change >0.5, pct.1 >0.25 and pct.2 <0.25. Ribosomal, mitochondrial, immunoglobulin and T cell receptor genes and tdTomato were excluded from RNA marker analyses.

### Gene signature scoring

Gene signature enrichment was assessed using AUCell. Gene sets representing resting Tregs, activated Tregs, effector Tregs, antigen-experienced Tregs and Treg identity were compiled from published signatures ^37, 38, 39, 40^. Genes not detected in the dataset were removed from each signature before scoring. For each cell, genes were ranked on the basis of normalized RNA expression, and AUCell area-under-the-curve scores were calculated using the top 5% of ranked genes. AUCell scores were added to Seurat metadata for downstream visualization and comparison.

For defined transcriptional programs, module scores were calculated using the Seurat AddModuleScore function. Gene sets representing Tfh, Th1, cytotoxic, quiescence/stability, proliferation and chronic activation programs were scored individually. Changes in module scores along Slingshot pseudotime were visualized using locally estimated scatterplot smoothing (LOESS).

### Transcriptional trajectory and pseudotime analysis

Transcriptional trajectory and pseudotime analyses of cTregs, eTregs and exTregs were performed using Monocle3^54^. The top 2,000 variable genes were used for trajectory analysis, and the Seurat WNN UMAP embedding was used as the reduced-dimensional representation. A principal graph was learned without partitioning or closed loops, with a minimum branch length of 25. Cells within the cTreg population at the beginning of the inferred trajectory were selected as root cells, and pseudotime was calculated using the Monocle3 “order_cells” function.

### Pseudotime-associated gene analysis

Pseudotime-associated genes were identified using tradeSeq ^44^. For analysis of the cTreg-to-eTreg-to-exTreg transition, generalized additive models were fitted to raw RNA counts for the top 2,000 variable genes using Monocle3 pseudotime and six knots. Dynamic genes were identified using the tradeSeq association test with Benjamini–Hochberg-adjusted P < 0.05. The top 150 dynamic genes ranked by adjusted P value were selected, and tradeSeq-fitted expression was calculated across 150 equally spaced pseudotime points. Gene-wise scaled expression profiles were grouped into six temporal modules by k-means clustering. Representative genes from each module were selected for visualization, and module scores were calculated using the Seurat AddModuleScore function and projected onto the WNN UMAP embedding.

For Shared and branch-specific dynamic gene analysis in exTreg subsets, tradeSeq was applied to each Slingshot lineage using the corresponding lineage-specific pseudotime. Analysis was restricted to the top 2,000 variable genes expressed in at least 5% of cells within each lineage. Generalized additive models were fitted using six knots, and pseudotime-associated genes were identified using the tradeSeq association test with Benjamini–Hochberg-adjusted P < 0.05. The top 100 dynamic genes for each lineage were selected, and fitted expression profiles across 100 pseudotime points were scaled by gene.

Dynamic genes identified by tradeSeq association testing were compared between paired lineages. For each lineage pair, the top dynamic genes were classified as shared genes (intersection) or branch-enriched genes (lineage-specific set difference). Gene expression along pseudotime was predicted from the joint multi-lineage tradeSeq model using predictSmooth. The endpoint of the shared developmental program was defined as the 95th percentile of pseudotime among cells in the final shared Slingshot cluster preceding lineage bifurcation (cluster 14 for the cytotoxic/Th1-cytotoxic trajectories and cluster 10 for the Tfh-like/proliferating trajectories). Smoothed expression profiles prior to the shared endpoint were averaged between paired lineages to visualize the common developmental program, whereas post-branch segments were displayed separately for each lineage. Expression values were row-wise Z-score normalized, ordered according to the pseudotime of maximal expression, and visualized as heatmaps. Whole-lineage pseudotime was normalized from 0 to 1, enabling direct comparison of the relative timing of shared and branch-specific transcriptional programs.

Transcription factor-associated genes were selected from lineage-specific dynamic genes, and their normalized expression along Slingshot pseudotime was visualized using LOESS.

### Slingshot analysis

Trajectory inference of exTreg states was performed using Slingshot^45^. Foxp3⁻tdTomato⁺ exTregs from the spleen were analyzed using WNN cluster identities and the WNN UMAP embedding generated in Seurat. Cluster 8 was specified as the starting cluster, and Slingshot was used to infer lineage connectivity and fit principal curves through the UMAP embedding. Lineage-specific pseudotime values and curve weights were extracted for each inferred lineage. For overall pseudotime analysis, each cell was assigned to the lineage with the highest curve weight, and the corresponding lineage-specific pseudotime was used as its dominant pseudotime. Cluster-centroid topology plots were generated by connecting cluster centroids according to the lineage structure inferred by Slingshot.

### TCR clonotype analysis

TCR repertoire analysis was performed using scRepertoire^55^. Cell Ranger V(D)J “filtered_contig_annotations.csv” files from the 15 samples were imported and combined using the “combineTCR” function. Clonotypes were defined using the strict clonotype assignment based on CDR3 amino acid sequence and V- and J-gene usage. TCR clonotype information was matched to the corresponding CD4^+ T cell Seurat metadata by cell barcode.

Clone size was calculated independently for each mouse by counting the number of cells assigned to each clonotype across tissues. Clones were classified as single (1 cell), small (2–5 cells), medium (6–20 cells) or large (21–100 cells). Cells without a valid strict clonotype assignment were considered unassigned. Clone-size classes were projected onto the WNN UMAP embedding and compared across tissues, Treg lineage populations and exTreg clusters.

Expanded clonotypes were defined as clonotypes represented by more than one cell. Clonotype numbers and clone-size distributions were quantified for cTregs, eTregs and exTregs and for individual exTreg clusters. Shared clonotypes between Treg lineage populations or exTreg clusters were identified by the presence of the same strict clonotype in two or more populations or clusters. Pairwise clonotype sharing between exTreg clusters was quantified as the number of overlapping clonotypes and visualized as a shared-clonotype matrix. A clonal network was constructed from pairwise exTreg cluster sharing, with nodes representing exTreg clusters, node size indicating the number of cells in each cluster and edge width proportional to the number of shared clonotypes.

For lineage-specific clonal analysis, exTreg clusters were grouped according to the lineage structures inferred by Slingshot. Within each lineage, shared expanded clonotypes were defined as clonotypes represented by more than one cell and detected in at least two clusters. Clonotypes were ranked by the number of clusters in which they were detected and by total cell number, and the most highly shared clonotypes were visualized across lineage-associated clusters. Clonotypes shared among exTreg states were similarly ranked by the number of exTreg clusters occupied and total cell number, and the top 30 shared clonotypes were visualized across exTreg clusters. Pairwise clonotype overlap between samples was assessed using the Jaccard index implemented in the scRepertoire “clonalOverlap” function.

### Statistical analysis

Statistical analyses were performed using GraphPad Prism (version 10.6.1) and R (version 4.4.3). Data are presented as mean ± SEM. unless otherwise indicated. Each data point represents an individual mouse or an independent cell-culture experiment, as specified in the figure legends. Comparisons between two independent groups were performed using an unpaired two-tailed Student’s t-test. Comparisons among three or more matched groups, including Treg populations or tissues derived from the same mice, were performed using repeated-measures one-way ANOVA followed by Tukey’s multiple-comparison test. Comparisons among three or more independent groups were performed using ordinary one-way ANOVA followed by Tukey’s multiple-comparison test.

For single-cell gene-signature analyses, distributions of AUCell scores between cell populations were compared using two-sided Wilcoxon rank-sum tests, with adjustment for multiple comparisons. Differential-expression analyses were performed using the Wilcoxon rank-sum test implemented in Seurat, and P values were adjusted using the Benjamini–Hochberg method. Pseudotime-associated genes were identified using the tradeSeq association test with Benjamini–Hochberg correction, as described above. Unless otherwise indicated, P < 0.05 was considered statistically significant. Exact sample sizes and statistical tests are provided in the corresponding figure legends. Significance is indicated as *P < 0.05, **P < 0.01, ***P < 0.001 and ****P < 0.0001; ns, not significant.

## Data availability

RNA-seq data have been uploaded to the NCBI GEO and are accessible under accession numbers GSExxxxxxx.

## Code availability

No new algorithms were generated for this study.

## Acknowledgements

We thank David Munn at Augusta University for providing Il6rafl/fl mice. We acknowledge the support and contributions of the Integrated Genomics Core Shared Resource at the Georgia Cancer Center, Augusta University (RRID: SCR_026483), and the Flow and Mass Cytometry Core Facility at the Georgia Cancer Center, Augusta University (RRID: SCR_025747).

## References

1. Wolf, D. & Ley, K. Immunity and inflammation in atherosclerosis. Circulation research 124, 315–327 (2019).

2. Kobiyama, K. & Ley, K. Atherosclerosis: a chronic inflammatory disease with an autoimmune component. Circulation research 123, 1118–1120 (2018).

3. Wang, Z. et al. Pairing of single-cell RNA analysis and T cell antigen receptor profiling indicates breakdown of T cell tolerance checkpoints in atherosclerosis. Nat Cardiovasc Res 2, 290–306 (2023).

4. Depuydt, M.A.C. et al. Single-cell T cell receptor sequencing of paired human atherosclerotic plaques and blood reveals autoimmune-like features of expanded effector T cells. Nature Cardiovascular Research 2, 112–125 (2023).

5. Wolf, D. et al. Pathogenic autoimmunity in atherosclerosis evolves from initially protective apolipoprotein B100–reactive CD4+ T-regulatory cells. Circulation 142, 1279–1293 (2020).

6. Kimura, T., Tse, K., Sette, A. & Ley, K. Vaccination to modulate atherosclerosis. Autoimmunity 48, 152–160 (2015).

7. Li, J. et al. CCR5+ T-bet+ FoxP3+ effector CD4 T cells drive atherosclerosis. Circulation research 118, 1540–1552 (2016).

8. Butcher, M.J. et al. Atherosclerosis-driven Treg plasticity results in formation of a dysfunctional subset of plastic IFNγ+ Th1/Tregs. Circulation research 119, 1190–1203 (2016).

9. Klingenberg, R. et al. Depletion of FOXP3+ regulatory T cells promotes hypercholesterolemia and atherosclerosis. J Clin Invest 123, 1323–1334 (2013).

10. Gotsman, I. et al. Impaired regulatory T-cell response and enhanced atherosclerosis in the absence of inducible costimulatory molecule. Circulation 114, 2047–2055 (2006).

11. Ait-Oufella, H. et al. Natural regulatory T cells control the development of atherosclerosis in mice. Nat Med 12, 178–180 (2006).

12. Mor, A. et al. Role of naturally occurring CD4+ CD25+ regulatory T cells in experimental atherosclerosis. Arterioscler Thromb Vasc Biol 27, 893–900 (2007).

13. Maganto-Garcia, E., Tarrio, M.L., Grabie, N., Bu, D.X. & Lichtman, A.H. Dynamic changes in regulatory T cells are linked to levels of diet-induced hypercholesterolemia. Circulation 124, 185–195 (2011).

14. Mallat, Z. et al. Protective role of interleukin-10 in atherosclerosis. Circ Res 85, e17–24 (1999).

15. Robertson, A.K. et al. Disruption of TGF-beta signaling in T cells accelerates atherosclerosis. J Clin Invest 112, 1342–1350 (2003).

16. Mallat, Z. et al. Inhibition of transforming growth factor-beta signaling accelerates atherosclerosis and induces an unstable plaque phenotype in mice. Circ Res 89, 930–934 (2001).

17. Zhou, X. et al. Instability of the transcription factor Foxp3 leads to the generation of pathogenic memory T cells in vivo. Nature immunology 10, 1000–1007 (2009).

18. Sakaguchi, S., Vignali, D.A., Rudensky, A.Y., Niec, R.E. & Waldmann, H. The plasticity and stability of regulatory T cells. Nat Rev Immunol 13, 461–467 (2013).

19. Sawant, D.V. & Vignali, D.A. Once a Treg, always a Treg? Immunol Rev 259, 173–191 (2014).

20. Smigiel, K.S., Srivastava, S., Stolley, J.M. & Campbell, D.J. Regulatory T-cell homeostasis: steady-state maintenance and modulation during inflammation. Immunol Rev 259, 40–59 (2014).

21. Hall, A.O. et al. The cytokines interleukin 27 and interferon-gamma promote distinct Treg cell populations required to limit infection-induced pathology. Immunity 37, 511–523 (2012).

22. Koch, M.A. et al. The transcription factor T-bet controls regulatory T cell homeostasis and function during type 1 inflammation. Nat Immunol 10, 595–602 (2009).

23. Koch, M.A. et al. T-bet(+) Treg cells undergo abortive Th1 cell differentiation due to impaired expression of IL-12 receptor beta2. Immunity 37, 501–510 (2012).

24. Levine, A.G. et al. Stability and function of regulatory T cells expressing the transcription factor T-bet. Nature 546, 421–425 (2017).

25. Bailey-Bucktrout, S.L. et al. Self-antigen-driven activation induces instability of regulatory T cells during an inflammatory autoimmune response. Immunity 39, 949–962 (2013).

26. Zhou, X. et al. Instability of the transcription factor Foxp3 leads to the generation of pathogenic memory T cells in vivo. Nat Immunol 10, 1000–1007 (2009).

27. Saxena, V., Lakhan, R., Iyyathurai, J. & Bromberg, J.S. Mechanisms of exTreg induction. European journal of immunology 51, 1956–1967 (2021).

28. Komatsu, N. et al. Pathogenic conversion of Foxp3+ T cells into TH17 cells in autoimmune arthritis. Nature medicine 20, 62–68 (2014).

29. Zhou, X., Bailey-Bucktrout, S., Jeker, L.T. & Bluestone, J.A. Plasticity of CD4+ FoxP3+ T cells. Current opinion in immunology 21, 281–285 (2009).

30. Oldenhove, G. et al. Decrease of Foxp3+ Treg cell number and acquisition of effector cell phenotype during lethal infection. Immunity 31, 772–786 (2009).

31. Vokaer, B. et al. Critical role of regulatory T cells in Th17-mediated minor antigen-disparate rejection. The Journal of immunology 185, 3417–3425 (2010).

32. Gaddis, D.E. et al. Apolipoprotein AI prevents regulatory to follicular helper T cell switching during atherosclerosis. Nature communications 9, 1–15 (2018).

33. Roy, P. et al. Loss of effector T(reg) signature in APOB-reactive CD4(+) T cells in patients with coronary artery disease. Nat Cardiovasc Res 4, 841–856 (2025).

34. Garg, G. et al. Blimp1 Prevents Methylation of Foxp3 and Loss of Regulatory T Cell Identity at Sites of Inflammation. Cell Rep 26, 1854–1868 e1855 (2019).

35. Hao, Y. et al. Integrated analysis of multimodal single-cell data. Cell 184, 3573–3587.e3529 (2021).

36. Oliaeimotlagh, M. et al. Automated denoising of CITE-seq data with ThresholdR. Cell Rep Methods 5, 101088 (2025).

37. Miragaia, R.J. et al. Single-Cell Transcriptomics of Regulatory T Cells Reveals Trajectories of Tissue Adaptation. Immunity 50, 493–504 e497 (2019).

38. Rosenblum, M.D., Way, S.S. & Abbas, A.K. Regulatory T cell memory. Nat Rev Immunol 16, 90–101 (2016).

39. Mijnheer, G. et al. Conserved human effector Treg cell transcriptomic and epigenetic signature in arthritic joint inflammation. Nat Commun 12, 2710 (2021).

40. Zemmour, D. et al. Single-cell gene expression reveals a landscape of regulatory T cell phenotypes shaped by the TCR. Nat Immunol 19, 291–301 (2018).

41. Huo, Y., Hafezi-Moghadam, A. & Ley, K. Role of vascular cell adhesion molecule-1 and fibronectin connecting segment-1 in monocyte rolling and adhesion on early atherosclerotic lesions. Circ Res 87, 153–159 (2000).

42. Ley, K. & Huo, Y. VCAM-1 is critical in atherosclerosis. J Clin Invest 107, 1209–1210 (2001).

43. Nakashima, Y., Raines, E.W., Plump, A.S., Breslow, J.L. & Ross, R. Upregulation of VCAM-1 and ICAM-1 at atherosclerosis-prone sites on the endothelium in the ApoE-deficient mouse. Arterioscler Thromb Vasc Biol 18, 842–851 (1998).

44. Van den Berge, K., et al. Trajectory-based differential expression analysis for single-cell sequencing data. Nat Commun 11, 1201 (2020).

45. Street, K. et al. Slingshot: cell lineage and pseudotime inference for single-cell transcriptomics. BMC Genomics 19, 477 (2018).

46. Harb, H. et al. A regulatory T cell Notch4-GDF15 axis licenses tissue inflammation in asthma. Nat Immunol 21, 1359–1370 (2020).

47. Huehn, J. et al. Developmental stage, phenotype, and migration distinguish naive- and effector/memory-like CD4+ regulatory T cells. J Exp Med 199, 303–313 (2004).

48. Kimura, T., Kobiyama, K. & Winkels, H. Regulatory CD4 (þ) T Cells Recognize MHC-II-Restricted Peptide Epitopes of Apolipoprotein B. Circulation 138, 1130–1143 (2018).

49. Lal, G. et al. Epigenetic Regulation of Foxp3 Expression in Regulatory T Cells by DNA Methylation1. The Journal of Immunology 182, 259–273 (2009).

50. Hua, J. et al. Pathological conversion of regulatory T cells is associated with loss of allotolerance. Sci Rep 8, 7059 (2018).

51. Helmin, K.A. et al. Maintenance DNA methylation is essential for regulatory T cell development and stability of suppressive function. J Clin Invest 130, 6571–6587 (2020).

52. Joudi, A.M. et al. Maintenance DNA methylation is required for induced Treg reparative function following viral pneumonia in mice. J Clin Invest 135 (2025).

53. Bankhead, P. et al. QuPath: Open source software for digital pathology image analysis. Sci Rep 7, 16878 (2017).

54. Trapnell, C. et al. The dynamics and regulators of cell fate decisions are revealed by pseudotemporal ordering of single cells. Nat Biotechnol 32, 381–386 (2014).

55. Borcherding, N., Bormann, N.L. & Kraus, G. scRepertoire: An R-based toolkit for single-cell immune receptor analysis. F1000Res 9, 47 (2020).

