## Supplementary Figures for "Atherosclerosis destabilizes regulatory T cells (Tregs) resulting in multiple families of exTregs"

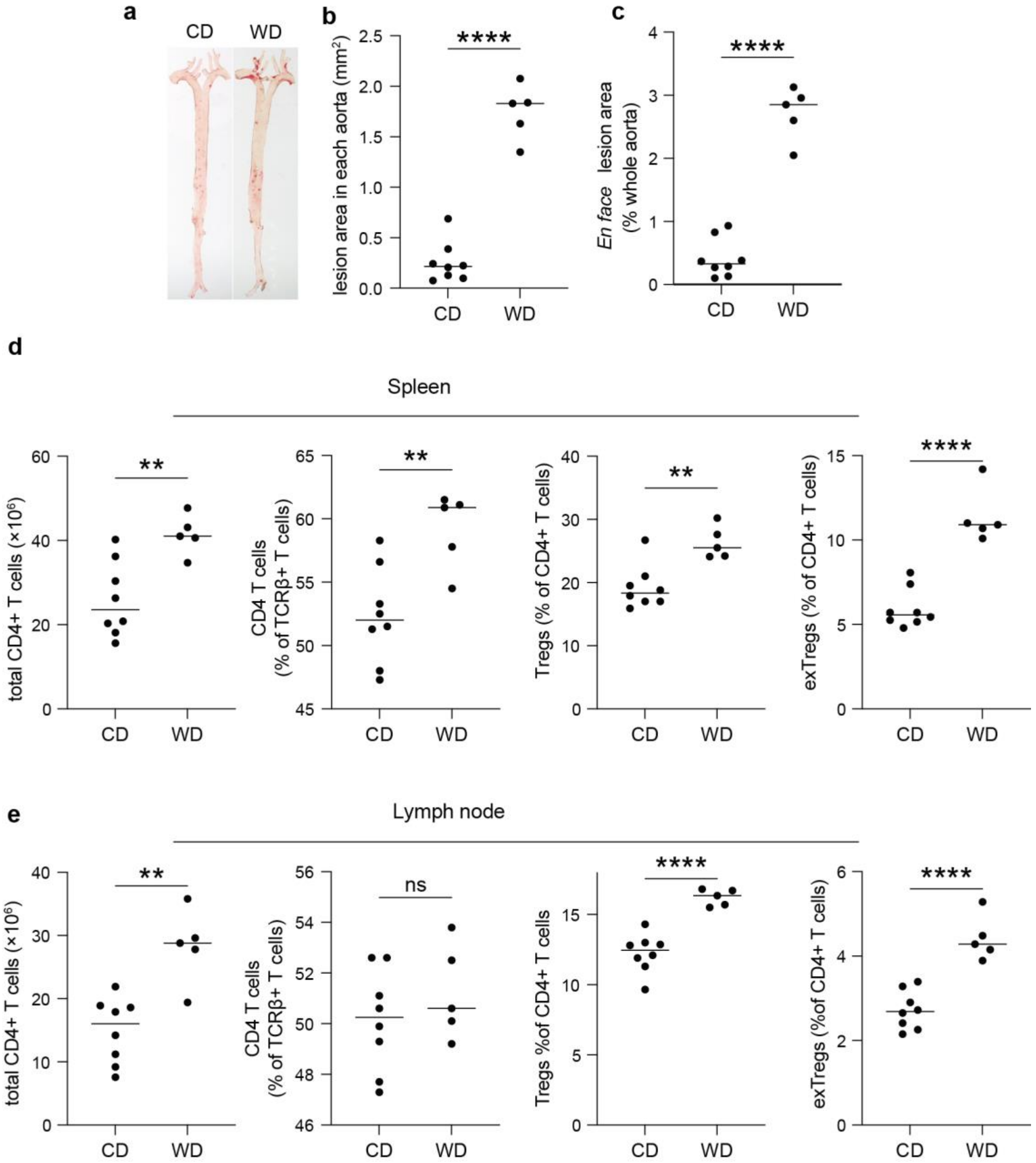
a
CD
WD
b
lesion area in each aorta (mm²)
****
CD
WD
c
En face lesion area (% whole aorta)
****
CD
WD
d
Spleen
total CD4+ T cells (×10⁶)
**
CD4 T cells (% of TCRβ+ T cells)
**
Tregs (% of CD4+ T cells)
**
exTregs (% of CD4+ T cells)
****
CD
WD
e
Lymph node
total CD4+ T cells (×10⁶)
**
CD4 T cells (% of TCRβ+ T cells)
ns
Tregs %of CD4+ T cells
****
exTregs (%of CD4+ T cells)
****
CD
WD

**Extended Data Fig. 1 | Establishment of the Foxp3 lineage-tracing atherosclerosis model. a,** Representative Sudan IV staining of whole aortas from chow diet (CD)- and Western diet (WD)-fed Foxp3 lineage-tracing $Apoe^{-/-}$ mice. **b and c,** Quantification of atherosclerotic lesion area per aorta (b) and en face lesion area as a percentage of total aortic surface area (c). **d,** Quantification of total CD4+ T-cell numbers, the frequency of CD4+ T cells among TCRβ+ cells, Tregs as a percentage of CD4+ T cells, and exTregs as a percentage of CD4+ T cells in the spleen. **e,** Quantification of total CD4+ T-cell numbers, the frequency of CD4+ T cells among TCRβ+ cells, Tregs as a percentage of CD4+ T cells, and exTregs as a percentage of CD4+ T cells in lymph nodes. Data are presented as mean ± SEM. Each dot represents one mouse. Statistical significance was determined using an unpaired two-tailed Student's *t*-test. *$P < 0.05$, **$P < 0.01$, ***$P < 0.001$, ****$P < 0.0001$.

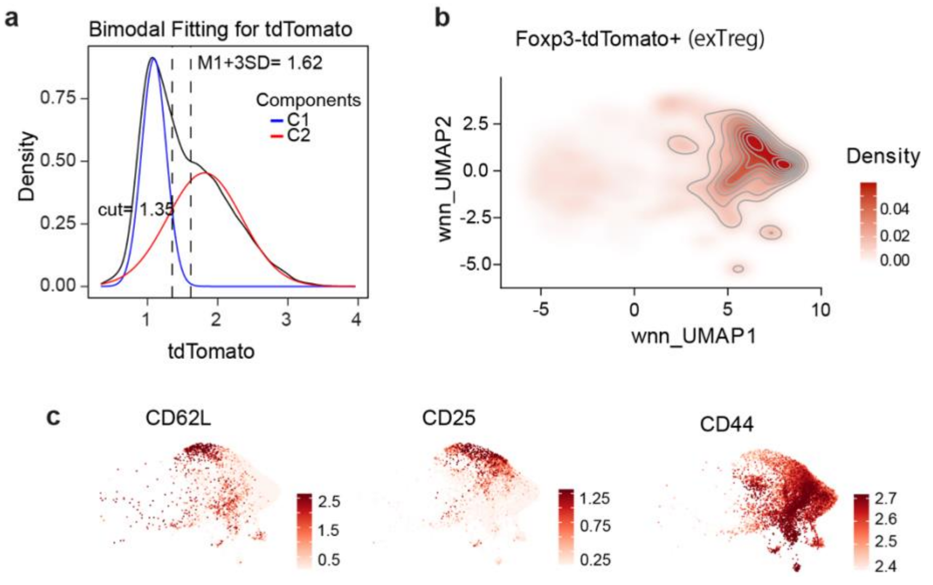


**Extended Data Fig. 2 | Annotation of lineage-traced Treg populations and validation of differentiation trajectory by diffusion map analysis. a,** ThresholdR analysis of tdTomato transcript expression showing bimodal distribution and objective determination of the cutoff separating tdTomato$^{low}$ and tdTomato$^{hi}$ populations. **b,** Density map of GFP-tdTomato+ cells projected onto the integrated WNN UMAP. Bona fide exTregs were defined as GFP−tdTomato+ cells enriched within the high-density region, whereas sparsely distributed GFP−tdTomato+ events outside this region were excluded as background. **c,** ADT feature maps showing representative surface marker expression (CD62L, CD25 and CD44) across the integrated single-cell dataset.

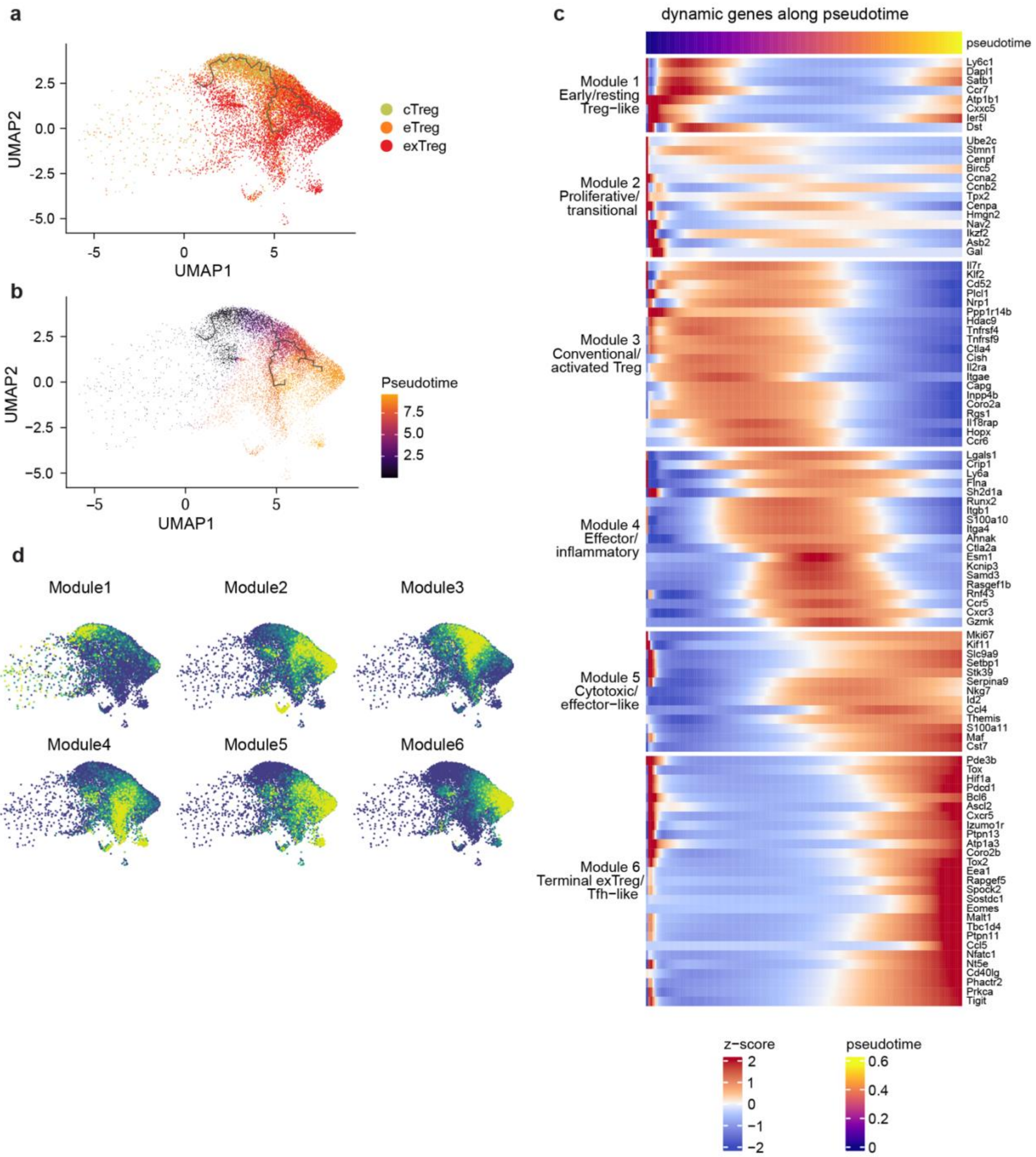
a
UMAP2
UMAP1
cTreg
eTreg
exTreg
b
UMAP2
UMAP1
Pseudotime
c
dynamic genes along pseudotime
pseudotime
Module 1 Early/resting Treg−like
Ly6c1
Dapl1
Satb1
Ccr7
Atp1b1
Cxxc5
Ier5l
Dst
Module 2 Proliferative/ transitional
Ube2c
Stmn1
Cenpf
Birc5
Ccna2
Ccnb2
Tpx2
Cenpa
Hmgn2
Nav2
Ikzf2
Asb2
Gal
Module 3 Conventional/ activated Treg
Il7r
Klf2
Cd52
Plcl1
Nrp1
Ppp1r14b
Hdac9
Tnfrsf4
Tnfrsf9
Ctla4
Cish
Il2ra
Itgae
Capg
Inpp4b
Coro2a
Rgs1
Il18rap
Hopx
Ccr6
Module 4 Effector/ inflammatory
Lgals1
Crip1
Ly6a
Flna
Sh2d1a
Runx2
Itgb1
S100a10
Itga4
Ahnak
Ctla2a
Esm1
Kcnip3
Samd3
Rasgef1b
Rnf43
Ccr5
Cxcr3
Gzmk
Module 5 Cytotoxic/ effector−like
Mki67
Kif11
Slc9a9
Setbp1
Stk39
Serpina9
Nkg7
Id2
Ccl4
Themis
S100a11
Maf
Cst7
Module 6 Terminal exTreg/ Tfh−like
Pde3b
Tox
Hif1a
Pdcd1
Bcl6
Ascl2
Cxcr5
Izumo1r
Ptpn13
Atp1a3
Coro2b
Tox2
Eea1
Rapgef5
Spock2
Sostdc1
Eomes
Malt1
Tbc1d4
Ptpn11
Ccl5
Nfatc1
Nt5e
Cd40lg
Phactr2
Prkca
Tigit
z−score
pseudotime
d
Module1
Module2
Module3
Module4
Module5
Module6

**Extended Data Fig. 3 | Monocle3 trajectory analysis identifies dynamic transcriptional programs during Treg destabilization. a,** Monocle3 trajectory reconstruction showing the inferred differentiation path from cTregs through eTregs to exTregs. **b,** Monocle3 pseudotime projected onto the inferred differentiation trajectory. **c,** Heatmap showing the top 100 pseudotime-dependent genes identified by tradeSeq based on the Monocle3 pseudotime trajectory and organized into six sequential transcriptional modules representing distinct stages of Treg differentiation. Modules correspond to early/resting Treg-like, proliferative/transitional, conventional/activated Treg, effector/inflammatory, cytotoxic/effector-like and terminal exTreg/Tfh-like transcriptional programs. **d,** Projection of the six transcriptional modules identified by tradeSeq onto the integrated single-cell landscape, illustrating their sequential activation during cTreg-to-exTreg differentiation.

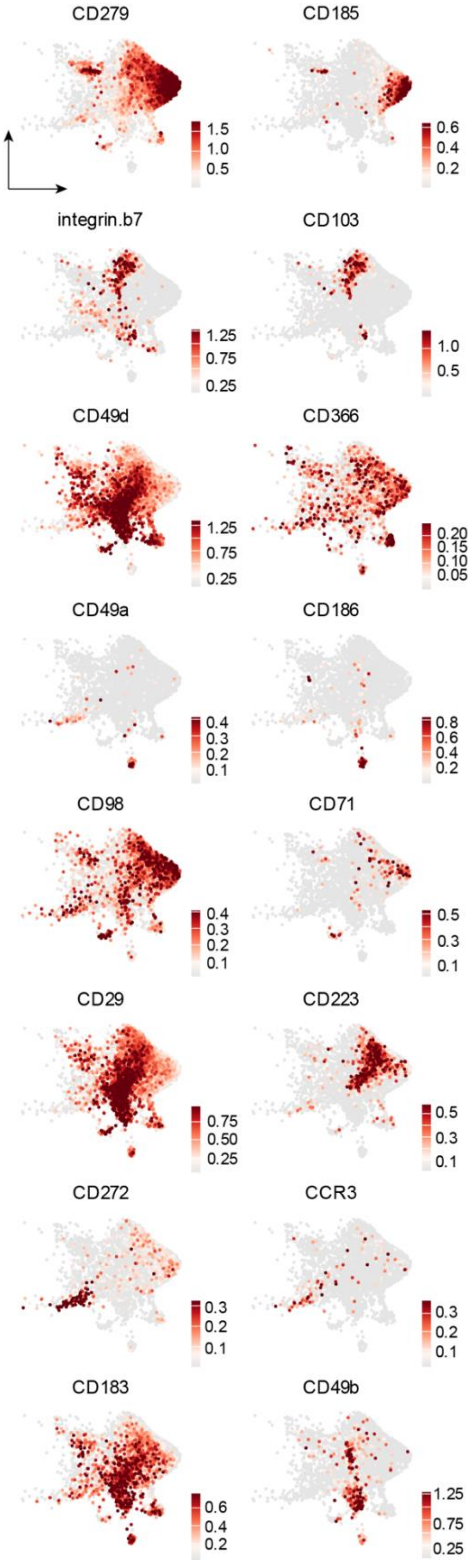
CD279
1.5
1.0
0.5
CD185
0.6
0.4
0.2
integrin.b7
1.25
0.75
0.25
CD103
1.0
0.5
CD49d
1.25
0.75
0.25
CD366
0.20
0.15
0.10
0.05
CD49a
0.4
0.3
0.2
0.1
CD186
0.8
0.6
0.4
0.2
CD98
0.4
0.3
0.2
0.1
CD71
0.5
0.3
0.1
CD29
0.75
0.50
0.25
CD223
0.5
0.3
0.1
CD272
0.3
0.2
0.1
CCR3
0.3
0.2
0.1
CD183
0.6
0.4
0.2
CD49b
1.25
0.75
0.25

**Extended Data Fig. 4 | Surface protein expression across exTreg states.** UMAP feature plots showing normalized antibody-derived tag (ADT) expression of CD279, CD185, CD103, integrin β7, CD49d, CD366, CD49a, CD186, CD98, CD71, CD29, CD223, CD272, CCR3, CD183 and CD49b across Foxp3-lineage Foxp3−tdTomato+ exTreg cells.

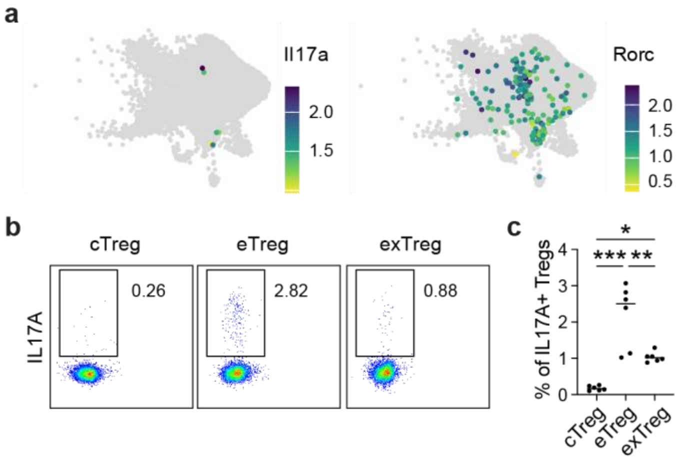


**Extended Data Fig. 5 | Limited acquisition of a Th17-like phenotype by exTregs. a,** UMAP feature plots showing expression of *Il17a* and *Rorc* in exTreg cells. **b,** Representative flow cytometry plots showing IL-17A expression in cTregs, eTregs and exTregs. **c,** Frequencies of IL-17A+ cells among cTregs, eTregs and exTregs. Each symbol represents an individual mouse ($n \geq 6$ mice per group). Data are presented as mean ± SEM. Statistical significance was determined by paired one-way ANOVA with multiple-comparison correction. *$P < 0.05$, **$P < 0.01$, ***$P < 0.001$, ****$P < 0.0001$, ns, not significant.

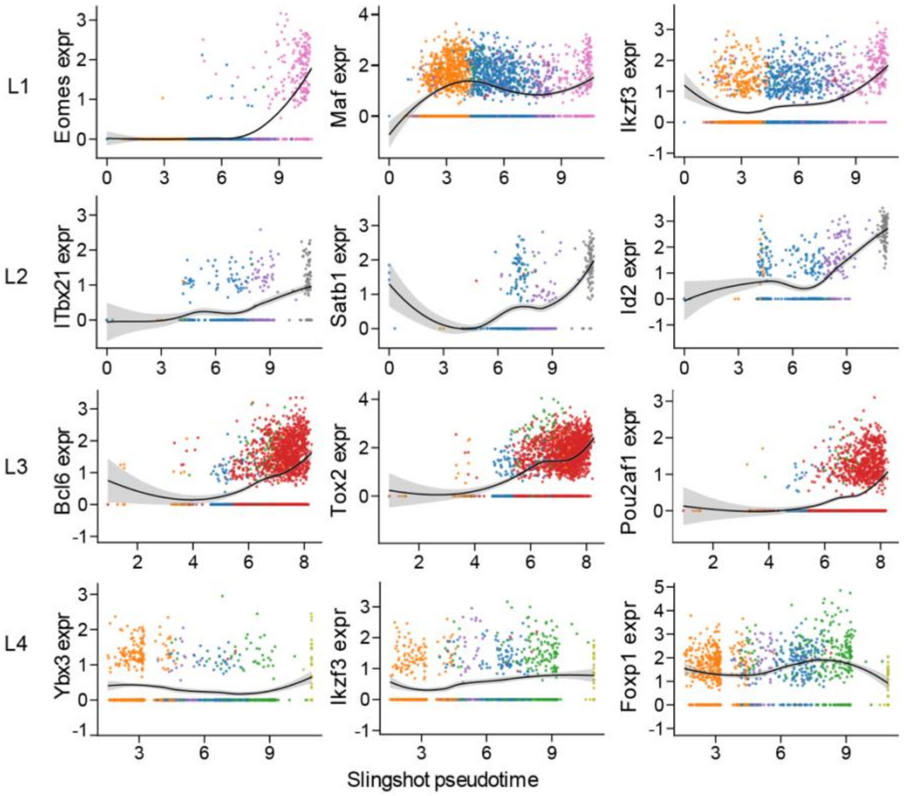


**Extended Data Fig. 6 | Lineage-specific transcription factor dynamics along exTreg differentiation trajectories.** Pseudotemporal expression dynamics of representative lineage-associated transcription factors selected from tradeSeq-identified dynamic genes. Expression is shown across Slingshot pseudotime for each exTreg lineage. Points represent individual cells and are colored by WNN cluster identity. Black lines indicate LOESS fits and shaded areas indicate 95% confidence intervals.

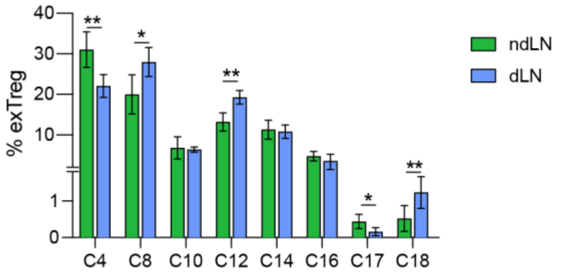


**Extended Data Fig. 7 | Composition of exTreg subsets in draining and non-draining lymph nodes.** Frequency of exTreg cells in each WNN cluster in ndLNs and dLNs. Each bar represents the mean across mice and error bars indicate s.e.m. Data are presented as mean ± SEM. n = 5 mice. Statistical significance was determined by paired one-way ANOVA followed by Tukey's multiple-comparison test. **P* < 0.05, ***P* < 0.01, ****P* < 0.001, *****P* < 0.0001, ns, not significant.

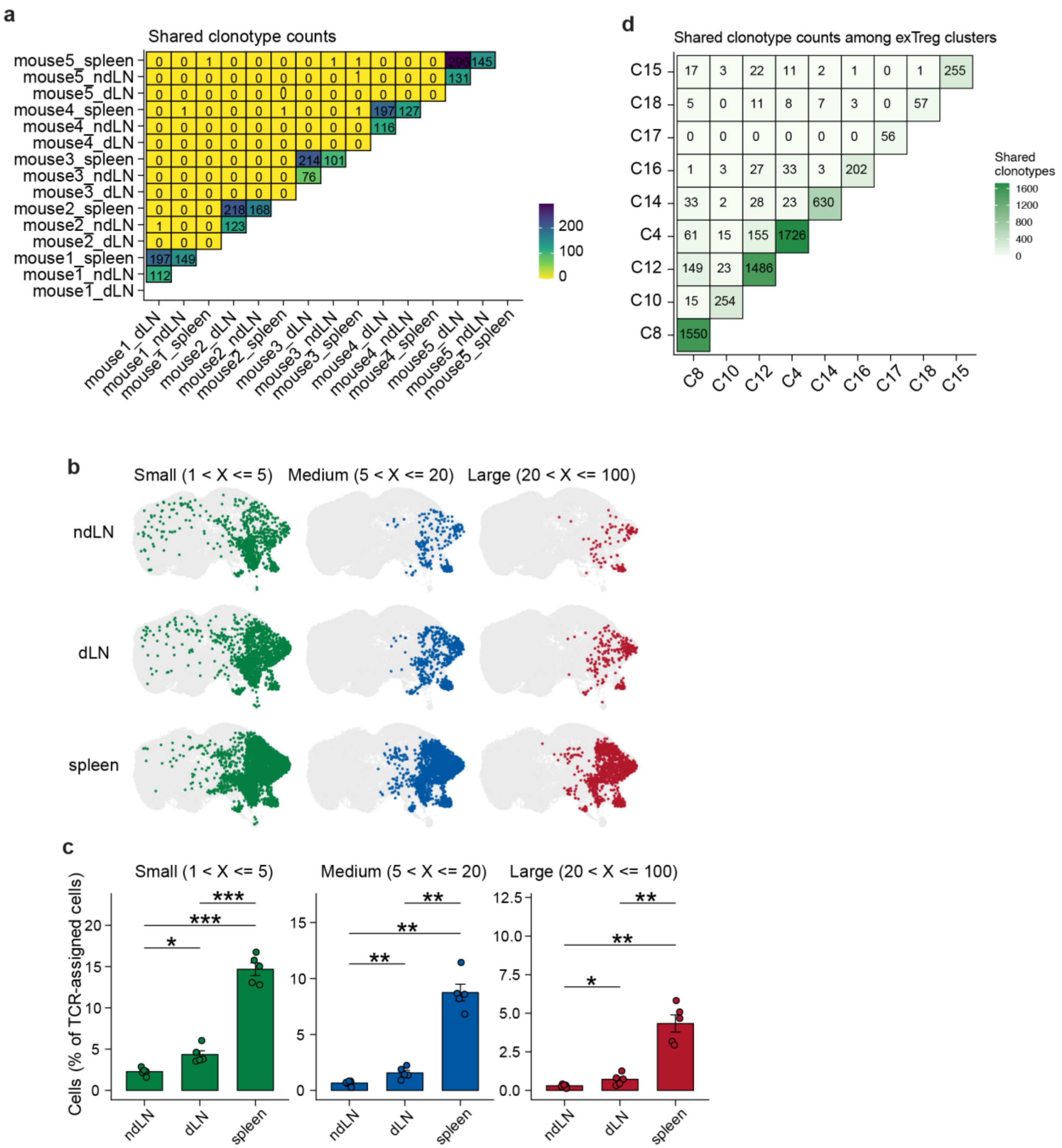

a
Shared clonotype counts
mouse5_spleen
mouse5_ndLN
mouse5_dLN
mouse4_spleen
mouse4_ndLN
mouse4_dLN
mouse3_spleen
mouse3_ndLN
mouse3_dLN
mouse2_spleen
mouse2_ndLN
mouse2_dLN
mouse1_spleen
mouse1_ndLN
mouse1_dLN
mouse1_dLN
mouse1_ndLN
mouse1_spleen
mouse2_dLN
mouse2_ndLN
mouse2_spleen
mouse3_dLN
mouse3_ndLN
mouse3_spleen
mouse4_dLN
mouse4_ndLN
mouse4_spleen
mouse5_dLN
mouse5_ndLN
mouse5_spleen
200
100
0
d
Shared clonotype counts among exTreg clusters
C15
C18
C17
C16
C14
C4
C12
C10
C8
Shared clonotypes
1600
1200
800
400
0
b
Small (1 < X <= 5)
Medium (5 < X <= 20)
Large (20 < X <= 100)
ndLN
dLN
spleen
c
Small (1 < X <= 5)
Medium (5 < X <= 20)
Large (20 < X <= 100)
Cells (% of TCR-assigned cells)
ndLN
dLN
spleen

**Extended Data Fig. 8 | Clonal expansion and clonotype sharing across Treg-lineage and exTreg states. a,** Pairwise numbers of shared clonotypes among ndLN, dLN and spleen samples from individual mice. Numbers and tile colors indicate the number of clonotypes shared between each pair of samples. **b,** WNN UMAPs showing the distribution of clonally expanded CD4+ T cells in ndLNs, dLNs and spleen. Clonotypes were classified according to clone size as small ($1 < x \leq 5$), medium ($5 < x \leq 20$) or large ($20 < x \leq 100$). Cells belonging to the indicated clone-size category are colored and other CD4+ T cells are shown in gray. **c,** Quantification of the clone-size distributions shown in (b). The frequencies of TCR-assigned CD4+ T cells belonging to small ($1 < x \leq 5$), medium ($5 < x \leq 20$) and large ($20 < x \leq 100$) clonotypes are shown for ndLNs, dLNs and spleen. Each dot represents one mouse; bars indicate the mean ± s.e.m. **d,** Pairwise numbers of shared clonotypes among exTreg clusters. Numbers and tile colors indicate the number of clonotypes shared between each pair of clusters. Diagonal values indicate the total number of clonotypes detected within each cluster. Clonotypes were defined using the scRepertoire strict clonotype definition, requiring identical paired TCRα and TCRβ CDR3 amino acid sequences and matching V and J gene usage. Data in **c** are presented as mean ± SEM. n = 5 mice. Statistical significance was determined by paired one-way ANOVA followed by Tukey's multiple-comparison test. *$P < 0.05$, **$P < 0.01$, ***$P < 0.001$, ****$P < 0.0001$, ns, not significant.
